# *In situ* 3D comparison of *Chlorella pyrenoidosa* with nuclear-irradiated mutagenic strains by using focused ion beam milling and cryo-electron tomography

**DOI:** 10.1101/2020.11.05.369421

**Authors:** Wangbiao Guo, Lingchong Feng, Zhenyi Wang, Jiansheng Guo, Donghyun Park, Brittany L. Carroll, Xing Zhang, Jun Liu, Jun Cheng

## Abstract

Microalgae are highly efficient photosynthetic organisms that hold enormous potential as sources of renewable energy. In particular, *Chlorella pyrenoidosa* displays a rapid growth rate, high tolerance to light, and high lipid content, making it especially valuable for applications such as flue gas CO_2_ fixation, biofuel production, and nutritional extracts. In order to unveil its full potential, it is necessary to characterize its subcellular architecture. Here, we achieved three-dimensional (3D) visualization of the architectures of *C. pyrenoidosa* cells, by combining focused ion beam scanning electron microscopy (FIB/SEM), cryo-FIB milling, and cryo-electron tomography (cryo-ET). These high-resolution images bring to light intricate features of intact organelles, including thylakoid membranes, pyrenoid, starch granules, mitochondria, nucleus, lipid droplets and vacuoles, as well as the fine architectures within the chloroplast, including the concave-convex pyrenoid, plastoglobules, thylakoid tips, and convergence zones. Significantly, comparative analysis of wild-type and nuclear-irradiated mutagenic strains determined that cell volume and surface area of mutant cells have increased substantially to 2.2-fold and 1.7-fold, respectively, consistent with up-regulation of the enzyme Rubisco and enhanced photosynthetic metabolic processes. Moreover, quantitative analysis established that the thylakoid membrane width in mutant cells increased to 1.3-fold, while the membrane gap decreased to 0.8-fold, possibly contributing to the higher biomass growth rate of mutant cells. Our work reveals the first 3D subcellular architectures of *C. pyrenoidosa* cell and provides a structural framework for unlocking the higher growth rate in microalgae relevant to a wide range of industrial applications.

## Introduction

The microalgae *Chlorella pyrenoidosa* is a spherical and unicellular eukaryotic organism belonging to class *Chlorophycea* and genus *Chlorella* that has evolved over more than 2 billion years (1). Among the tens of thousands of microalgal species, *Chlorella* is one of the most unique species because of its rapid reproduction ability. It owes to its unique materials *Chlorella* Growth Factor, a nucleotide peptide complex enriched with vitamins, minerals, and carbohydrates, which makes it in a quadripartion-division reproduction method. In theory, one *Chlorella* cell can reproduce 100 million cells within 10 days, nearly equally to the speed of nuclear fission (2). Besides, *C. pyrenoidosa* is increasingly used for industrial and biomedical applications, such as CO_2_ flue gas fixation (3, 4), biodiesel production (5, 6), high-value nutritional extracts (7–10), and anti-inflammatory products (1, 11). Moreover, nuclear radiation (12–14) is commonly used in full-scale bioreactors (15) to realize specified functions, such as increasing removal of pollutants from wastewater (16), higher CO_2_ concentration fixation (17), and increased biodiesel production (18). Importantly, the organelles within the cell function collaboratively to support its rapid growth and sufficient metabolic activities. Understanding the subcellular architectures of microalgae cells, including the structural basis underlying how mutations due to nuclear radiation increase biomass growth rate, stands to unlock their full benefits as renewable energy sources.

Most existing knowledge of this subcellular architecture has been acquired by Scanning Electronic Microscopy (SEM) and Transmission Electron Microscopy (TEM) (19–21). In particular, TEM was used to visualize *C. pyrenoidosa* organelles, such as the chloroplast, nucleus and mitochondria (22). The 2D TEM shows that the chloroplast is cup-shaped, and contains a pyrenoid, plenty of starch plates, and the double thylakoid membranes (23). However, conventional TEM sample preparation (e.g. fixation, dehydration, infiltration, plastic embedding, and sectioning with the ultramicrotome) is unable to reveal native cellular features in sufficient detail. Furthermore, 2D TEM micrograph is inadequate to reveal organelles in three dimensions (3D). Therefore, it is important to provide an overall and comprehensive 3D view of the intact *C. pyrenoidosa* cell at a high resolution in order to gain detailed insights into its subcellular world.

With the development of Focused Ion Beam Scanning Electron Microscopy (FIB-SEM) and its biological applications, it is now possible to image intact cells in 3D by FIB-SEM (24, 25). Furthermore, cryo-FIB milling coupled with cryo-electron tomography (cryo-ET) provides *in situ* snapshots of frozen-hydrated cells at sufficient resolution (26) to further view the subcellular architecture within the organelles. Therefore, the combination of FIB-SEM and cryo-FIB/cryo-ET provides a unique avenue to study the subcellular architecture of the *C. pyrenoidosa* cell.

Here, we combined various advanced imaging techniques to visualize the subcellular architecture of wild-type and nuclear-irradiated *C. pyrenoidosa* cells. Qualitative analysis by FIB-SEM revealed that the volume and surface area of organelles were increased substantially. In addition, cryo-FIB milling and cryo-ET imaging exposed the intricate features inside chloroplasts. Our findings provide the first direct structural evidence to explain the higher biomass growth rate of nuclear-irradiated *C. pyrenoidosa* cell.

## Results

### Mutation of *C. pyrenoidosa*

To select mutants with faster growth rate, wild-type *C. pyrenoidosa* suspensions were radiated under the ^*60*^*Co γ*rays with the radiation dose of 500 GY and dose-rate of 15 GY/min (27, 28). Through this process, the natural evolution of cells is dramatically accelerated, as most of the genetic sequences of microalgae cells are broken down and reorganized. Among hundreds of mutants (15), one mutant *C. pyrenoidosa* (MCP) has a biomass growth rate 24% higher than that of wild type strain (WCP) at the cultivation time of 72 h (Fig. S1A). To compare the WCP and MCP in detail, we developed two independent workflows to visualize the cells (Fig. S1A).

### 3D visualization of wild and mutant *C. pyrenoidosa* cells by FIB-SEM

To preserve the high contrast ultrastructure of *C. pyrenoidosa* cells, we use high-pressure freezing together with subsequent freeze substitution and resin embedding to prepare WCP and MCP specimens. The resin embedded specimen blocks were then imaged by FIB-SEM (Fig. S1B-D). To obtain the complete dataset, 1,199 and 1,481 frames of the image stacks were acquired for the WCP and MCP cells, respectively (Fig. S1E-F, Video 1, 2). We selected four intact cells of WCP and MCP, respectively (Fig. 1C, D, W1 – W4 and M1 – M4) for 3D visualization and detailed analyses. 3D structures of the *C. pyrenoidosa* cells provide distinct morphologies of organelles: cell wall (gray), nucleus (cyan), pyrenoid (red), mitochondria (orange), thylakoid (green), vacuoles (yellow), lipid droplets (tan), starch grains (blue) and chloroplast (purple) (Video 3, Video 4). The overall shape and organization of the organelles in WCP (W1 in Fig. 1) and MCP (M1 in Fig. 1) are similar (Fig. 2A, B), however there are some noticeable differences: WCP is 25.0±4.2 μm^3^ in cell volume, while MCP is considerably increased 2.2-fold to 54.0±29.8 μm^3^ in cell volume (Fig. 2C, Table S1, S2). Compared to the surface area (82.8±14.7 μm^2^) of WCP, MCP has increased 1.7-fold to 139±50.7 μm^2^ (Fig. 2E, Table S4).

**Figure 1.**
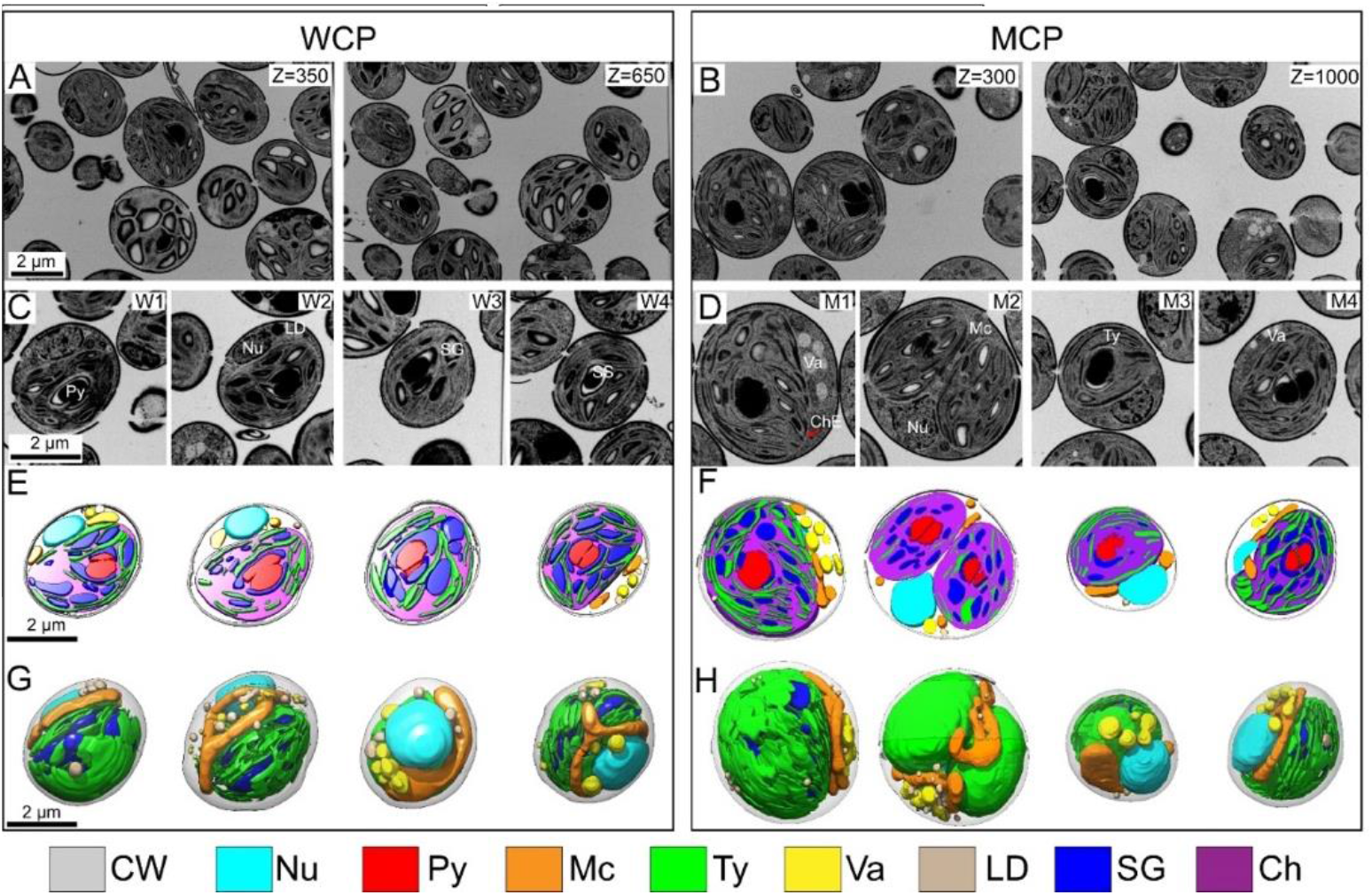
Comparison of intact 3D architecture of wild (WCP)and mutant (MCP) *C. pyrenoidosa* cells by FIB-SEM. (A, B) The raw FIB-SEM images of WCP and MCP at the different *z* axial planes. (C, D) The central plane of four evenly distributed intact cells of WCP and MCP, respectively. (E, F) The corresponding segmentation section of the same MCP and WCP cells with the lamella depth of 400 nm. (G, H) The corresponding intact 3D segmentation architecture of the same WCP and MCP cells. Colors: gray-cell wall (CW), cyan-nucleus (Nu), red-pyrenoid (Py), orange-mitochondria (Mc), green-thylakoids (Ty), yellow-vacuoles (Va), tan-lipid droplets (LD), blue-starch granules (SG), purple-chloroplast (Ch).

**Figure 2.**
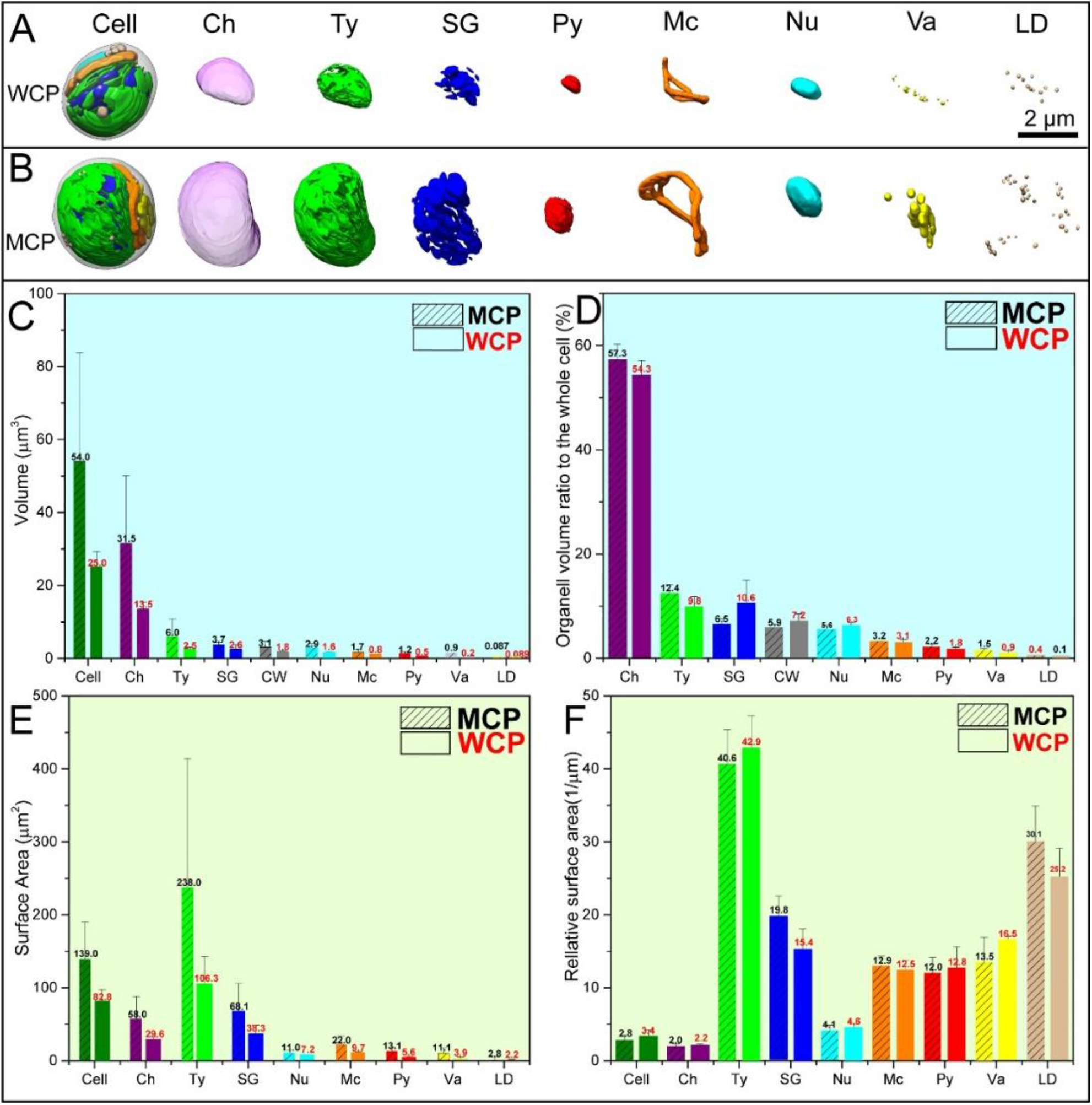
Comparison of organelles architecture, and their corresponding volume and surface area of wild (WCP) and mutant (MCP) *C. pyrenoidosa* cells. (A, B) Organelles comparison of WCP and MCP by FIB-SEM (W1 and M1 in Fig. 2). The chloroplast envelope (ChE), thylakoid (Ty), starch granule (SG), pyrenoid (Py), mitochondria (Mc), nucleus(Nu), vacuole (Va) and lipid droplets (LD) were compared. (C) The average volume of the whole cell and their organelles. (D) The corresponding volume ratio of organelle to the whole cell. (E) The average surface area of the whole cell and their organelles. (F) The corresponding relative surface area of cell and organelles, which is the ratio of surface area to volume.

### Comparative analyses of organelles in *C. pyrenoidosa*

The architecture of chloroplast in the WCP is ellipsoidal-like, the average volume is 13.5±1.9 μm^3^, which makes-up 54.3% of the cell volume (Fig. 2E, F, Table S5). This result is similar with the previous observation (25). The chloroplast consists of thylakoid membrane, pyrenoid, starch granules and chloroplast matrix, and is encompassed by the chloroplast envelope (Video 5). Inside the chloroplast, the volume of thylakoid and starch granules occupies 18.0% and 19.3% of the chloroplast, respectively. The pyrenoid and chloroplast matrix occupy 3.4% and 59.3%, respectively (Table S6). The flaky-like thylakoid membrane is prolonged from one-side of the chloroplast to another side and eventually converged to the rim of the chloroplast. The average volume of thylakoid is 2.5±0.6 μm^3^, which makes-up 9.8% of the whole cell. The surface area of thylakoid is 106.3±36.7 μm^2^, correspondingly the relative surface area is 42.9±4.4 μm^−1^. A twin-shaped pyrenoid is localized in the center of chloroplast, which is covered by the concave-convex starch sheaths (Fig. S2E). Part of the thylakoid membranes are penetrated through the middle gap of the pyrenoid (Fig. 1D, M4). Starch granules are widely synthesized in the chloroplast matrix and wrapped by the thylakoids (Both of starch sheath and starch granules were colored as blue). The average volume of all the starch granules are 2.6±1.1 μm^3^, and the pyrenoid is 0.5±0.1 μm^3^ (Fig. 2C, Table S1), which makes-up 10.6% and 1.8% (Fig. 2D, Table S3), respectively.

A mitochondria loop is usually existed between nucleus and chloroplast to guarantee their ATP supply, and it is prolonged along one side of chloroplast to another side (Fig. S2F, Video 6). The mitochondria in the WCP is continuous, amorphous and loop-shaped (Fig. S2A-D). Besides, an ellipsoidal-like nucleus is usually located at the side of the chloroplast, where is far away from the central position of the cell. The average volume of nucleus and mitochondria is 1.6±0.2 μm^3^ and 0.8±0.1 μm^3^, respectively (Fig. 2C, Table S1), which makes-up 6.3% and 3.1% of the whole cell (Fig. 2D, Table S3). The surface area of nucleus and mitochondria is 7.2±0.5 μm^2^ and 9.7±2.3 μm^2^, respectively (Fig. 2E, Table S4), while the relative surface area of nucleus and mitochondria is 4.6±0.4 μm^−1^ and 12.5±1.3 μm^−1^, respectively (Fig. 2F, Table S5).

The electron-dense lipid droplets (Fig. 1C, W2) and electron-light vacuoles (Fig. 1D, M1) in the WCP are scattered among the nucleus and mitochondria alternatively and randomly in the cytoplasm. The architecture of single lipid droplet and vacuole is ellipsoidal-like or ball-like. The volume of single vacuole is higher than that of lipid droplet. The single lipid droplet and vacuole is concentrated and assembled, respectively, which are located along the periplasmic space of mitochondria and nucleus (Fig. 2A, B, Va and LD). The volume of lipid droplets and vacuoles is 0.089±0.023 μm^3^ and 0.2±0.1 μm^3^, respectively (Fig. 2C, Table S1), which makes-up 0.4% and 0.9% of the whole cell (Fig. 2D, Table S3). The surface area of lipid droplets and vacuoles is 2.2±0.3 μm^2^ and 3.9±1.7 μm^2^, respectively (Fig. 2E, Table S4).

Overall, the volume size of chloroplast, thylakoid, starch grains, nucleus, mitochondria, pyrenoid and vacuole in MCP have increased to 2.33 fold, 2.45 fold, 1.41 fold, 1.81 fold, 2.18 fold, 2.52 fold, and 4.04 fold, which jointly resulted in the increase of cell volume size by 2.2 fold (Fig. 2C, Table S1). Furthermore, the surface area of chloroplast, thylakoid, starch grains, nucleus, mitochondria, pyrenoid, vacuole and lipid droplets in MCP have increased to 9.37 fold, 2.24 fold, 1.78 fold, 1.52 fold, 2.27 fold, 2.31 fold, 2.84 fold and 1.28 fold (Fig. 2E, Table S4), which resulted in the increase of cell surface area by 1.7 fold.

### Chloroplast architecture revealed by cryo-FIB milling and cryo-ET

FIB-SEM enables visualization of the intact *C. pyrenoidosa* cells, providing morphological differences between WCP and MCP. To obtain detailed information of the *C. pyrenoidosa* cells, cryo-FIB and cryo-ET were used to achieve higher resolution. Under cryo-ET, we were able to see the double layers of the cell wall, cell membrane and the scattered ribosomes in the cytoplasm (Fig. S3F). The double membrane of the chloroplast envelope is easily visualized under cryo-ET while we can only visualize an electron-light line under FIB-SEM (Fig. S3F, Fig. 2D). Besides, the converge of starch granules by thylakoid membrane is clearly visualized under cryo-ET (Fig. S3G). The shape of starch sheaths and thylakoid membrane penetrating through the pyrenoid are visualized under cryo-ET while the pyrenoid and starch sheaths under FIB-SEM are only black features (Fig. S3D, H). Moreover, we have visualized the double membrane of mitochondria under cryo-ET, while we can only visualize the outline of mitochondria under FIB-SEM (Fig. S3E, I). Therefore, it is possible to explore more details of the *C. pyrenoidosa* at higher resolution by cryo-ET imaging (Fig. S1G-K).

We have milled a lamella of WCP showing the flaky-like thylakoid membrane penetrating through the concave-convex starch sheaths (Fig. 3). The starch sheath is covered by a single membrane, which may be the boundary of starch, blocking the leakage of Rubisco complexes. Outside the starch sheath, the thylakoid membranes distribute in an annular shape to cover the starch sheath. Moreover, there exist two thylakoid sheet penetrating through the starch sheath at the entry of starch sheath (Fig. 3E, F), which connects the two sides of the thylakoid membranes (up and down). The thylakoid membrane penetrating through the starch sheath is flaky-like, and the concave-convex starch sheaths is continuous. The thylakoid membrane can only penetrate through the middle entry gate of the starch sheath (Fig. 3H, I).

**Figure 3.**
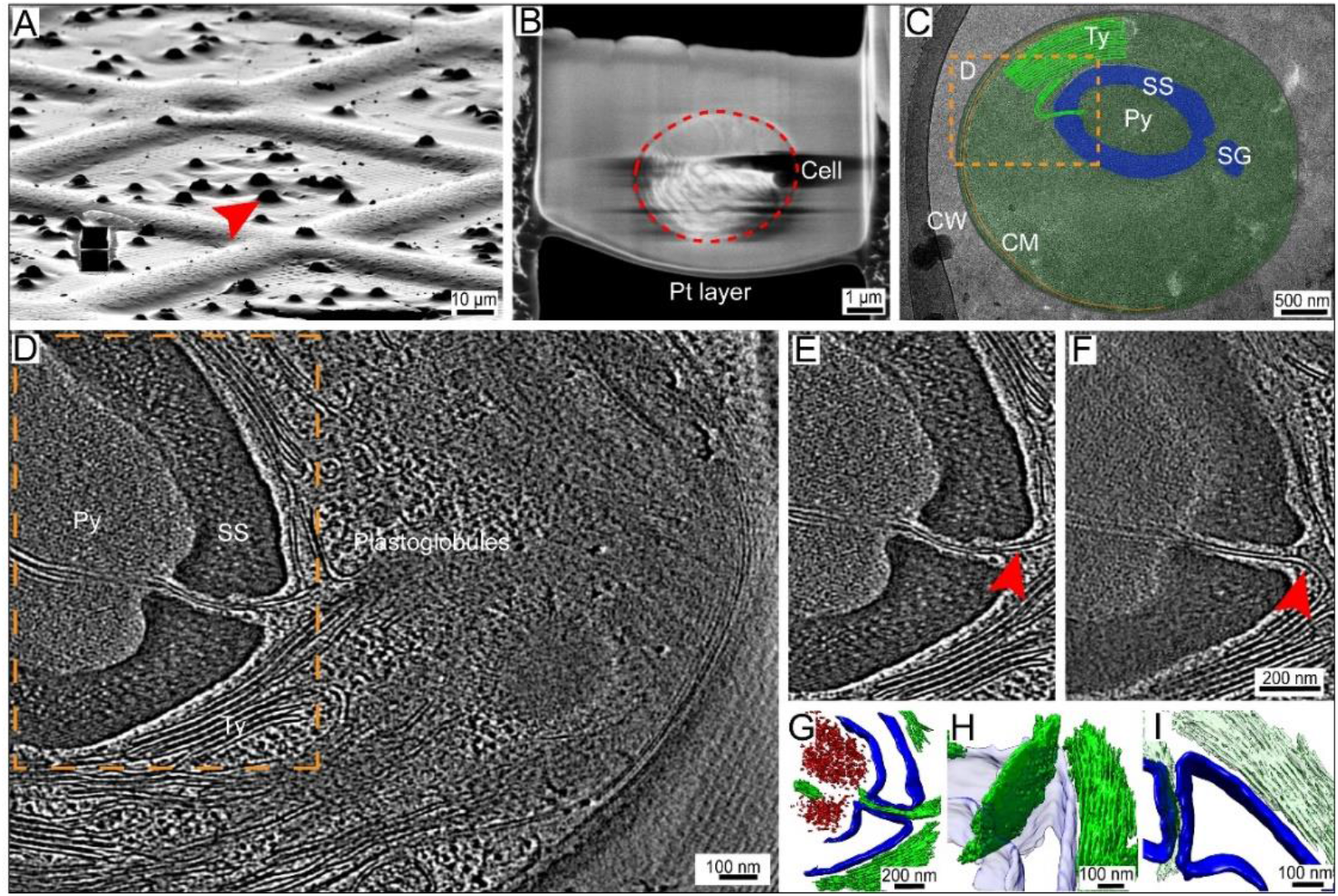
A cryo-FIB milled lamella of a wild *C. pyrenoidosa* cell showing the thylakoid membrane penetrating through the pyrenoid. (A) The cell imaged with the FIB before milling at the tilt angle of 18°. (B) Top-view of SEM image of the milled lamella from the same cell shown in A. The milling direction is from bottom to top. The red-dotted circle is the cell section covered in the lamella. The Pt layer on the bottom of lamella is to protect the sample from damage by the Ga+ ion beam. The final depth of lamella is 200 nm. (C) TEM high-defocus montaged overview of the same lamella in (B), which showed the position of pyrenoid. (D) A typical tomographic slice acquired at the position of orange-dotted box in (C). The pyrenoid (Py) is filled with Rubisco complexes and covered by two-halves starch sheath (SS). The starch sheath is a single layer electron-dense structure. The plastoglobules were visualized among the thylakoid membrane. (E-F) Two typical slices of thylakoid membranes (up and down) came across the pyrenoid, connecting the light-dependent reaction with light-independent reaction. (G-I) The corresponding 3D segmentation model. (G) The 3D segmentation model at the direction of (E). Colors: blue-starch sheath, red-pyrenoid, green-thylakoids. (H) Rotation of (G) showed the flaky-like structure of thylakoid membranes coming across the starch sheath instead of tubules-like. (I) Rotation of (G) showed the top view of starch sheath, which is strictly divided into two halves by the thylakoid membranes.

The circular connection channel, which connects the inner envelope of chloroplast (IE) with thylakoid membrane, was observed in our tomograms (Fig. 4G, J, Fig. S4). The circular connection channel is part of IE, and appears at the rim of the chloroplast. The IE extends its membrane to shorten the distance with thylakoid membrane, but it does not merge with the thylakoid membrane, and the IE is still a closed membrane system. 3D segmentation shows that the circular connection channel is a tubular-like architecture instead of flaky-like, and the length is around 170 nm.

**Figure 4.**
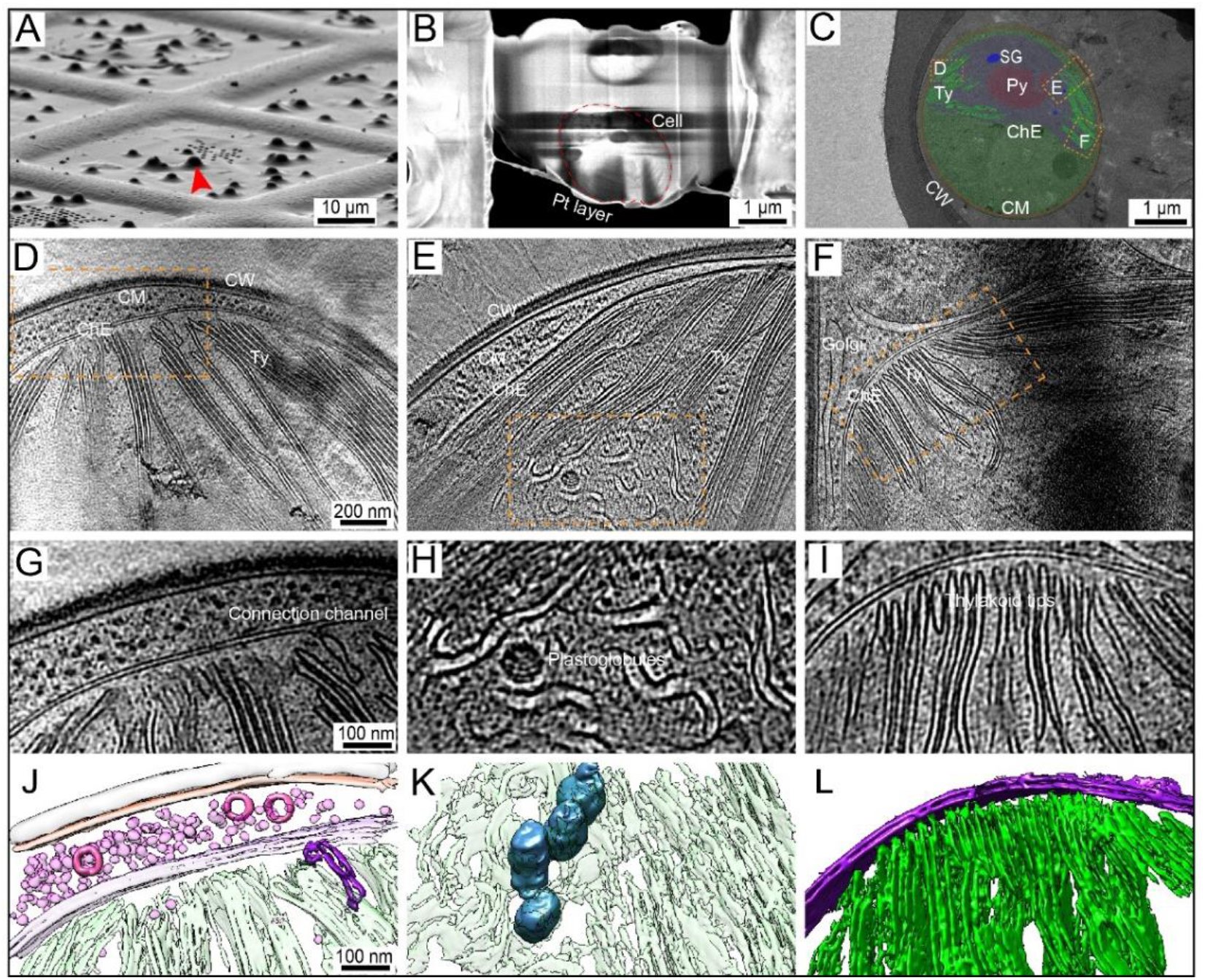
A cryo-FIB milled lamella of a mutant *C. pyrenoidosa* cell showing the detailed thylakoid membrane features. (A) The cell imaged with the FIB before milling at the tilt angle of 18°. (B) Top-view of SEM image of the milled lamella from the same cell shown in A. The milling direction is from bottom to top. The red-dotted circle is the cell section covered in the lamella. The final depth of lamella is around 200 nm. (C) TEM high-defocus montaged overview of the same lamella in (B). The chloroplast envelope (ChE), thylakoid membrane (Ty), pyrenoid (Py) and starch granules (SG) were visualized. (D, G, J) Detailed visualization of a circular connection channel connecting the chloroplast inner envelope and thylakoid membrane. (D) A typical tomographic slice imaged at the position of orange-dotted box in (C) showing the circular connection channel. (G) Zoom in visualization of orange-dotted box in (D). (J) The corresponding 3D segmentation model of (G). (E, H, K) Direct visualization of plastoglobules among the thylakoid membranes. (E) A typical tomographic slice imaged at the position of orange-dotted box in (C) where the warped thylakoid membrane and plastoglobules were identified. (H) Zoom in visualization of orange-dotted box in (E). The double thylakoid membrane is warped to cover the plastoglobules. (K) The corresponding 3D segmentation model of (H). (F, I, L) Thylakoid tips convergence zone was founded at the rim of chloroplast. (F) A typical tomographic slice imaged at the position of orange-dotted box in (C) showing the thylakoid tips. (I) Zoom in visualization of orange-dotted box in (F). (L) The corresponding 3D segmentation model of (I). Colors: purple-chloroplast envelope, steel blue-plastoglobules, magenta-ribosomes, pink-vesicles, green-thylakoids, orange red-cell membrane, gray-cell wall.

The single and grouped plastoglobules clusters were visualized in the chloroplast matrix, and were covered by the thylakoid membranes with a high curvature (Fig. 4H, K, Fig. S5). To cover the plastoglobules, the double membrane of thylakoids was extended and decomposed to single membrane. The thickness of thylakoid membrane was increased at the position of plastoglobules appear, where close to the starch granules. Moreover, similar to starch granules, a single membrane layer is existed outside the plastoglobules to avoid the leakage of nutrient materials (Fig. S5). 3D segmentation shows that the plastoglobules are cylindrical-like, with the diameter of 76±10 nm and length of 87±14 nm.

Thylakoid tips, which formed a thylakoid tips converge zone, were visualized (Fig. 4I, L). We have visualized this architecture both by the FIB-SEM and cryo-ET data, which shows it a common phenomenon. The arrangement of thylakoids at the rim of chloroplast is parallel or near parallel. 3D segmentation shows that nearly all the thylakoid membranes turn towards the chloroplast envelope. The density of thylakoid membranes at this zone is dense. The Golgi apparatus appears to cover the chloroplast envelope (Fig. 4I).

Moreover, we have visualized the microtubule-like architectures in the chloroplast matrix (see the forest green in Fig. S7). Different from the parallel-organized membrane of thylakoids, theses thylakoid membranes were warped and formed as microtubule-like (Fig. S6E). 3D segmentation shows that they are continuous and exist among the thylakoid membranes, the average diameter of these microtubules is 161±22 nm. The distance of microtubules perpendicular to the closest thylakoids is 64±6 nm. Besides, the width of thylakoid double membranes was enlarged at this area and the cross section of some warped thylakoid membranes was turned to shuttle-shaped, with the average maximum width of 47±18 nm.

### Thylakoid membrane comparison of wild and mutant *C. pyrenoidosa* cell

The thylakoid membrane width of MCP has increased to 1.33-fold while the membrane gap has decreased to 0.8-fold (Fig. 5). We have measured 7 tomograms of WCP and 9 tomograms of MCP, and totally 100 datasets were measured for WCP and MCP, respectively (Table S7). The thylakoid membrane, which close to the chloroplast matrix, was defined as outer membrane (OM), which close to the thylakoid stroma, was defined as inner membrane (IM). The distance between inner membrane and outer membrane was defined as membrane width, and between inner membrane and inner membrane was defined as membrane gap. The distance between IM and OM was defined as membrane width, and between IM and IM was defined as membrane gap (Fig. 5G). Results show that the membrane width of WCP and MCP is 13.0±2.3 nm and 17.3±2.2 nm, respectively. The membrane gap of WCP and MCP is 10.4±1.3 nm and 8.2±1.0 nm, respectively (Table S7). Therefore, the thylakoid membrane width of MCP has increased to 1.3-fold while the membrane gap has decreased to 0.8-fold.

**Figure 5.**
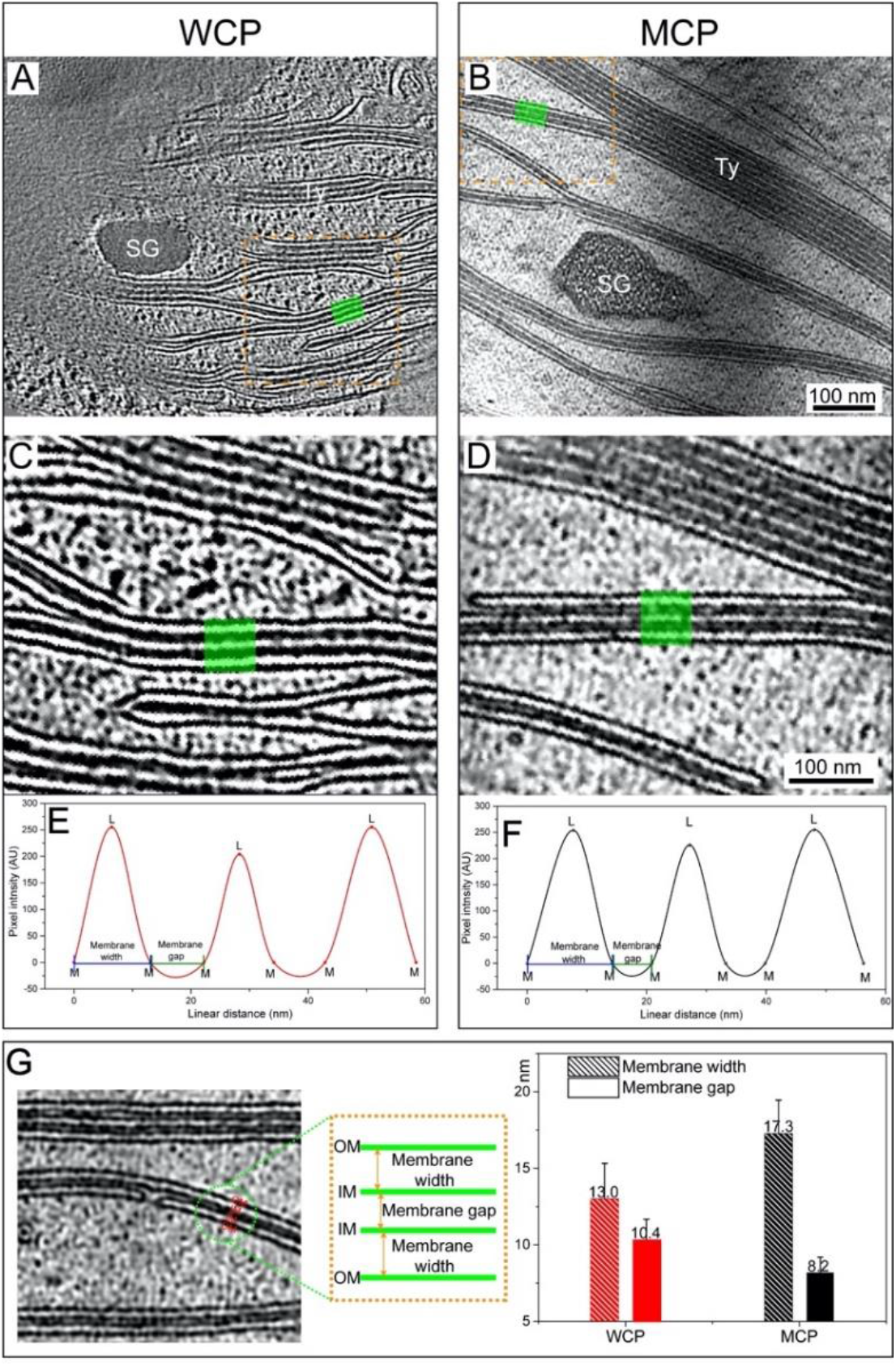
Thylakoid membrane comparison of wild (WCP)and mutant (MCP) *C. pyrenoidosa* cells. (A, B) The typical thylakoid membrane stacks visualized in WCP and MCP cells, respectively. (C, D) Zoom in visualization of orange-dotted box in (A), (B), respectively. (E, F) The pixel intensity along the green shadows in (C, D) from top to bottom. The linear distance is the real distance of green shadows which was measured with Image J (NIH, USA). M: thylakoid membrane, L: thylakoid lumen. AU: arbitrary units. Typically, the thylakoid is the double-membrane structure and two or more groups of double-membranes are assembled together. Combined with (G), the thylakoid membrane, which close to the chloroplast matrix, was defined as outer membrane (OM), which close to the thylakoid stroma, was defined as inner membrane (IM). The distance between inner membrane and outer membrane was defined as membrane width, and between inner membrane and inner membrane was defined as membrane gap. (G) The comparison of average membrane width and membrane gap between MCP and WCP cells. In order to eliminate the errors, we have summarized 100 membrane stacks of MCP and WCP cells (see Table S7).

### Enhanced physiological and metabolism process in chloroplast of mutant *C. pyrenoidosa* cell

FIB-SEM and cryo-FIB/cryo-ET have revealed major morphologic differences between WCP and MCP. To further understand these differences, we conducted the proteomics and metabolomics analysis (Fig. S8–S11). We found that the proteins, involved in glyoxylate and dicarboxylate metabolism, carbon fixation in photosynthetic organisms, and porphyrin and chlorophyll metabolism, were up-regulated to 1.4-fold, 1.4-fold and 1.3-fold in MCP (Fig. S8). In contrast, other proteins within pyruvate metabolism and biosynthesis of unsaturated fatty acids were down-regulated to 0.81-fold and 0.75-fold in MCP, respectively. Evidently, compared to WCP, photosynthesis in MCP was enhanced while subsequent generation of secondary metabolites such as fatty acid biosynthesis was weakened. These results are consistent with major morphologic differences between WCP and MCP.

## Discussion

### 3D visualization by FIB-SEM provide quantitative results of wild and mutant *C. pyrenoidosa*

We have obtained the first 3D view of intact *C. pyrenoidosa* cell by FIB-SEM. Our results showed that the largest organelle in the wild *C. pyrenoidosa* cell is the chloroplast, which makes-up around 54% of the cell volume. Mitochondria, as the ATP supplier, is spatially closely connected the chloroplast with nucleus to support their ATP requirements. So the chloroplast and nucleus were separated by the looped mitochondrion (Fig. S2F). The mitochondria architecture is stretched from one side of the chloroplast to another side, there may be an ATP transport channel between chloroplast and mitochondria. It was proposed that the NADPH produced at the chloroplast was transferred to the cytosol and oxidized by the mitochondrial electron chain (29). Our cryo-ET data showed the outer membrane of chloroplast and mitochondria were parallel, while the obvious connection channel was not identified (Fig. S3I). The spatial distribution of *C. pyrenoidosa* cell is compact and ordered, providing basis for its efficient biological activities.

Both starch granules and lipid droplets are energy-storage compounds which are in an inversely proportional relationship. The component of lipid droplet is triacylglycerol (TAG), which is an energy-rich compound to protect the cells against photoinhibition and to be utilized for membrane rearrangement in cells in case of extreme environmental conditions (30). It is suggested that the volume ratio of starch granules and lipid droplets in the *C. pyrenoidosa* cell is dependent to the microalgae cell division rate. At the initial inoculum and fast growth stage, the cell concentration is low and cell division rate is fast. At this stage, the larger starch granules are the prevalent storage materials all over the chloroplast, whereas the small lipid droplets occasionally appeared in the chloroplast or cytosol (31). Our data were acquired at the cultivation time of 72 hours, when the cell is still under the “rapid growth period”. The nutrients are sufficient and cell dividing is the most prevalent physiological activities. Therefore, starch granules were intensively visualized while only limited lipid droplets were identified in the cytosol. The appearance of vesicles may be to do with the triacylglycerol transport towards lipid droplets (31). So the lipid droplets and vesicles were simultaneously observed in the cytosol in our FIB-SEM data (Fig.1).

The cell wall of our high-pressure frozen cell samples were broken (see Fig. 1), and we noticed the similar phenomenon from other works (25). The reason may be that the external cryoprotectant (1-hexadecene) was not spread out evenly to the cell samples, thus the samples were suffered directly from the instant high pressure (over 2000 bars), which may destroy the cell wall of the sample. But the inside cellular features were not affected. The main component of the cell wall is cellulose and gelatin, which is kind of polysaccharide substance, with a function of maintaining the cell morphology and increasing the cell mechanical strength. The molecule composition resulted in the rigidity of the cell (32). The instant high pressure would disrupt the surface tension and resilience within the cell wall, thus causing the crack in the surface of the cell wall. Golgi and ER were not displayed in our 3D segmentation models, as they are too fuzzy to be detected clearly in our FIB-SEM image.

### Proteomics and metabolomics analyses of wild and mutant *C. pyrenoidosa*

The proteomics and metabolomics results are consistent with our imaging results. In carbon metabolism process, proteins related to Calvin cycle, especially a series of RubisCO were universally overexpressed in MCP (Fig. S8A) (33). In addition to the light reaction and Calvin cycle of photosynthesis, other metabolic processes which occurred or partially occurred in the chloroplast were also generally enhanced and resulting in an increase of related metabolites, including pyrimidines and pyrimidine derivatives, purines and purine derivatives, phenylacetamides, carbonyl compounds, carbohydrates and carbohydrate conjugates (Fig. S9B). Besides, although the processes related to amino acid metabolism showed a differentiation of enhancement and weakness. Among them, the lysine degradation process mainly happened in chloroplast was enhanced because of the enhancement of chloroplast. In summary, the basis for these enhanced chloroplast metabolic processes may be the general overexpression of chloroplast ribosomal proteins (Fig. S8A), which implied the enhancement of protein translation and structure formation in chloroplast (34). As a result, the volume of chloroplast and various structures (thylakoids, pyrenoid, etc.) in chloroplast were increased in MCP.

Oxidative phosphorylation dominated processes in mitochondria were generally weakened in MCP. It was possibly because cytochrome *C* synthesis, which was responsible for mitochondrial respiration, was a competitive pathway with chlorophyll synthesis (35, 36). Or possibly because the TCA cycle, a precursor to oxidative phosphorylation, required the product of glycolysis as raw material (37), but starch decomposition was inhibited and provided less pyruvates for TCA cycle and following respiration in MCP. Besides, metabolism processes which partially occurred in mitochondria such as alanine, aspartate and glutamate metabolism, valine, leucine and isoleucine degradation were also weakened (Fig. S10B). However, in mitochondria of MCP, the buffer of malate and oxaloacetate was sufficiently mobilized to shift the reaction (R00342) balance towards the consumption of malate and synthesis of oxaloacetate to provide additional NADH for oxidative phosphorylation (Fig. S11A). The mitochondrial respiration would be inhibited by a high cytosolic ATP/ADP ratio (29). Therefore, although the physiological functions in mitochondria were generally weakened by the decreased volume ratio of mitochondria in MCP, MCP still tried to provide more substrates for the normal operation of oxidative phosphorylation to supply enough ATP for cell photosynthetic growth. The second evidence was that another substrate succinic acid in oxidative phosphorylation was significantly reduced in MCP (Fig. S11A), which showed that in the absence of a weakened TCA cycle, mitochondria tried to use limited substrates for normal respiratory processes. But even so, the respiration was still weakened because the protein structure for TCA cycle and oxidative phosphorylation was reduced because of the reduced volume ratio of mitochondria. Therefore, the active transport process ABC transporters that consumed ATP would be also weakened (Fig. S10A).

The abundance of actin located in nucleus was significantly decreased in MCP (Fig. S8A), which may affect transcription, chromatin organization and nuclear structure (38). Because of the damage of actin in nucleus, the volume ratio and relative surface area of nucleus in MCP were all decreased (Fig.2D, F). As a result, pyrimidine nucleosides and purine nucleosides that made up genetic materials were reduced (Fig. S9A-B). Besides, cysteine and methionine metabolism partially occurred in nucleus also weakened (Fig. S10B). In addition, there was a significant reduction in the metabolites (D-Sedoheptulose 7-phosphate) associated with the pentose phosphate pathway, which was the previous process of genetic material synthesis (Fig. S11B). Therefore, a general damage to physiological functions in nucleus were happened because of the nuclear structural damage caused by downregulated expression of actin. As a result, the massive synthesis of genetic material that is essential for cell division was blocked in MCP, which led to the cell enlargement (from 25.0 μm^3^ to 54.0 μm^3^) caused by cell division inhibition (39).

In contrast to enhanced photosynthetic carbon fixation, the whole carbon metabolism process showed an inhibition inclination (Fig. S10A), thus the metabolism process of generating starch granules was enhanced while lipid droplets inhibited. An evidence was the abundance of proteins associated with starch decomposition and subsequent glucose decomposition was generally decreased (Fig. S8A). Then the proteins associated with biosynthesis of unsaturated fatty acids were following decreased (Fig. S8B). The other evidence showed in metabolites: Some carbohydrates and carbohydrate conjugates increased, but all types of lipids and lipid-like molecules generally decreased (Fig. S9A-B). This meant that inorganic carbon in MCP, once fixed as starch, tended to exist as starch rather than continue to be metabolized as lipid or other secondary metabolites. This result is consistent with our FIB-SEM data.

### New insights into the chloroplast of *C. pyrenoidosa* by cryo-FIB/cryo-ET

Different from the tubule-like architecture of thylakoid membrane penetrating through the pyrenoid in *Chlamydomonas reinhardtii* cell (40), in *C. pyrenoidosa* cell, it is flaky-like. Besides, the thylakoid membranes can only penetrate through the middle gate of the pyrenoid matrix (Fig. 3, Fig. S2E). It is investigated that the pyrenoid is a phase-separated, liquid-like organelle (41–43) and wherein most global CO_2_ fixation occurs in the vicinity of Rubisco (ribulose-1,5-bisphosphate carboxylase/oxygenase) through CO_2_-concentrating mechanisms (CCM). In the theory of CCM, the microalgae cell can utilize the HCO_3_^−^ as the extra carbon supply to eliminate the low CO_2_ concentration in the atmosphere. The inorganic carbon HCO_3_^−^ is imported into the cell by the bicarbonate transporters LCI1 and HLA3 with a form of protein complex on the cell membrane. Alternatively, CO_2_ molecule may also enter the cell and be converted to HCO_3_^−^ possibly by LCIB. Then the cytoplasmic HCO_3_^−^ is mediated by LCIA on the chloroplast envelope and into the thylakoid lumen. The function of thylakoid membrane penetrating through the pyrenoid is to provide a channel for the diffusion of HCO_3_^−^ to the pyrenoid, where it is converted to CO_2_ by CAH3 in turn (44). Inside the pyrenoid, it is filled with the complex of EPYC1 binding with four Rubiscos. In our work, we visualized the dotted substances, which may be the complex of EPYC1 and Rubiscos. We have visualized the flaky-like thylakoid membrane architecture. It has wider contact area with light, which may be beneficial to the light absorption ability. So the material and energy exchange mechanism between thylakoid-membrane light-reaction and pyrenoid dark-reaction in C. *pyrenoidosa* cell may be different from that of in *Chlamydomonas reinhardtii* cell. The function of starch sheath is to prevent the leakage of CO_2_ from the pyrenoid and to minimize the oxygen exposure in the pyrenoid matrix (45, 46). Therefore, the pyrenoid, starch sheath, and thylakoid membranes jointly consist of the *Ci* transport and utilization system, connecting the light-dependent and light-independent reactions. The flaky-like thylakoid membrane and concave-convex pyrenoid are beneficial to the photosynthesis of *C. pyrenoidosa*.

The circular connection channel, which connects the IE and thylakoid membrane, may be to do with the material and energy transmission between chloroplast envelope and thylakoid membrane (Fig. 4J). It is reported that the orchestrated regulation of IE is effective to the thylakoid membrane structure and volume regulation in higher plants. The osmotically active IE would be invaginated from the stromal side towards thylakoid lumen when there is a net influx of ions (such as H_2_O, Cl^−^, Ca^2+^, Mg^2+^, et al). The cation channels of IE could assist control chloroplast volume (47, 48). As a high-sufficient photoautotrophic organism, the photosynthetic efficiency of *C. pyrenoidosa* cell is high, so the existence of circular connection channel may provide an ion channel accelerating the ion and ATP exchange rate. It gave the evidence that the chloroplast bridges the thylakoid via invaginations from the IE instead of via shuttle of soluble transfer proteins or via the cargo protein (49). The occurrence of plastoglobules were associated with the thylakoid membranes with a high curvature (Fig. 4H, K, Fig. S4). They are functioned as lipid biosynthesis and storage subcompartments of thylakoid membranes, and are physically covered by the thylakoid membranes via a half-lipid bilayer. The biochemical composition of plastoglobules includes prenyl quinones, vitamin K1, free fatty acid, carotenoids, and chlorophylls (50–52). The plastoglobules related proteins include plastoglobulins family, chloroplast and chromoplast metabolic proteins (51). Our data have visualized the single or grouped plastoglobules in the thylakoid membranes where with a high curvature. Different from higher plants, the plastoglobules in *C. pyrenoidosa* cell appears at the position close to starch granules. It is likely that there exists a conversion mechanism between lipids and starch to regulate the energy storage method. The thylakoid tips were converged to the rim of chloroplast, forming a thylakoid tip convergence zone (Fig. 4I, L). It illustrated that the thylakoid double membrane is a closed membrane system. The formation of thylakoid convergence zone may be concerned with CURT1 protein family (40, 53). There is evidence that the PratA-defined membrane (PDMs) protein were identified at the thylakoid tip convergence zone in cyanobacteria, which involved in the synthesis and assembly of photosynthetic compartments. While in eukaryotic organism, there is no direct evidence to show the existence of PDMs protein (49). The microtubule-like architecture is a kind of thylakoid membrane form in the chloroplast matrix (see the forest green in Fig. S7). It has been reported in the cyanobacteria (54) and algae (55), which hypothesized with the function of bridging the inner envelope and thylakoid membranes. While our tomography data is obvious different. The formation of this kind of microtubule-like architecture may be the over-expression of chloroplast-construction related proteins (Fig. S8).

Moreover, the increased thylakoid membrane width and decreased membrane gap of MCP (Fig. 5) may be to do with the enhanced carbon metabolism process. As the Calvin cycle related proteins, especially Rubisco enzyme, were universally overexpressed in MCP (Fig. S8A) (33). Correspondingly, the chlorophyll synthesis proteins of light-independent protochlorophyllide reductase were increased, which made positive effect on light reaction of carbon fixation. To load increased light harvesting pigments for providing stronger photosynthetic light reaction, a structural transformation happened in chloroplast to meet the requirements of the photosynthetic reaction by increased Rubisco. Therefore, the surface area of chloroplast and thylakoid increased to provide wider light harvesting area. The thickness of thylakoid membrane became widened and the gap between membranes became narrow, which made the transport distance between the membranes shorter to transport photosynthetic electrons and coenzymes quicker. Thus the transport efficiency was accelerated, and the communication efficiency between light reactions and dark reactions was generally improved.

In summary, we combined FIB-SEM and cryo-FIB/cryo-ET imaging to visualize the subcellular 3D architecture of the *C. pyrenoidosa* cell for the first time and reveal major morphological changes in a nuclear-irradiated mutagenic strain in which the cell volume and growth rate have increased substantially.

## Materials and Methods

### Cell culture

The wild-type *Chlorella pyrenoidosa* strain (FACHB-9) was acquired from Freshwater Algae Culture Collection at the Institute of Hydrobiology, Chinese Academy of Sciences, China. The nuclear-irradiation was conducted in the Institute of Nuclear-Agricultural Sciences, Zhejiang University, China. The detailed methods were introduced by the previous work (28). Both of mutant and wild *C. pyrenoidosa* cells were grown in BG 11 medium (56) with constant light (~23200 lux) and aerated with CO_2_ gas (0.1 vvm) at 27 °C room temperature.

### High pressure freezing (HPF) and freeze substitution (FS)

The HPF and FS were conducted with previous proposed protocol (57). The centrifuged microalgae samples (ALLEGRAX-15R, 1000 rpm, 3mins) were dipped into external cryoprotectant (1-hexadecene) and carefully loaded into sample carriers (0.1 mm depth, Type A, Leica). The lid (Type B, Leica) was dipped in 1-hexadecene and placed on top of the sample carrier. And the samples were frozen using an EM ICE (Leica) HPF device (Fig. 1 (B)). After which the samples were substituted in an EM AFS2 (Leica). The sample chamber was filled with ethyl alcohol to keep the temperature stability. The specimen carrier was transferred into a cap containing 1% uranyl acetate and 2% OsO_4_ in the liquid nitrogen, usually every cap stored two carriers (Fig. 1C). After FS, samples were washed with acetone (2 times for 30min each) at 4°C, followed by acetone (2 times for 15 min each) at room temperature. Then the acetone was substituted with a mixture of acetone: epoxy resin=7:3 and incubated for 8 h. After that, the former solution (acetone: epoxy resin=7:3) was replaced by the mixture of acetone: epoxy resin=3:7 and incubated for 12 h. Afterwards, it was changed to pure epoxy resin incubating for 24 h. Followed by exchanging with pure epoxy resin for another 24 h. At the end, the Polymerize epoxy resin was heated at 60 °C in an oven for 2 days.

### Focused ion beam milling combined with scanning electron microscopy (FIB-SEM)

The high-pressure frozen samples were embedded as resin blocks, which were trimmed using Leica EM trimmer carefully until the surface of black tissue in the block was seen. After finding the area of interest, the resin blocks were imaged under a dual beam scanning electron microscopy (Thermo Fisher, FIB Helios G3 UC) ((Fig. S1D). The data collection procedure was operated in the serial-surface view mode with the slice thickness of 8 nm and current of 0.79 nA at 30 keV. Each serial face was then imaged with a 2.5kV acceleration voltage and a current of 0.2 nA in backscatter mode with an ICD detector. The image store resolution was set to 6144 × 4096 pixels with a dwell time of 2 μs per pixel and 2.44 nm per pixel. And 1199 and 1481 slices were obtained of WCP and MCP cells. The horizontal field width of images of WCP and MCP is 18.6 and 16.3 μm, respectively.

### Plunge freezing

Usually 5 μL of liquid culture (OD_680_ ~ 0.3) was pipetted onto holey carbon-coated copper grids (R2/1, 200 mesh, Quantifoil Micro Tools, Jena, Germany). The grids were blotted from the reverse side and immediately plunged into the liquid ethane at liquid nitrogen temperature atmosphere using a Vitrobot. The Vitrobot was set to room temperature, humidity 90%, blot force 10, and blot time 7 s (Fig. S1G). The grids were stored in sealed boxes in liquid nitrogen until used (40).

### Cryo-focused ion beam milling (Cryo-FIB)

Cryo-FIB milling was performed as described in previous work (58). Before milling, the chamber vacuum should be checked to lower than 4 ×10^−4^ Pa. Then heat exchanger was inserted to dewar, meanwhile flow of cryo-stage and cryo-shield were set to 7~8 L/min and waiting their temperature to stabilized below −180 °C for 10 mins. The temperature of cryo-stage and cryo-shield were always kept under −170 °C during the whole milling process. After which, the plunge-frozen grids were mounted into the dual-beam (FIB-SEM) microscope (Aquilos, Thermal Fisher, USA) via an autoloader loading station. At this step, the samples should always under liquid nitrogen or vacuum atmosphere. During cryo-FIB operation, the initial samples state was evaluated (ice, thickness, et al) under low magnification (100~200 x, voltage 2 kv and dwell time 300 ns). Then launching the Maps software to take a grid image at low magnification (100~200 x). Adjusting the magnification, voltage and dwell time until the grid bars being clearly (typically, 800~1000 x, 5 kv and 3 μs) so as to take a montage image, where the lamella sites can be chosen. Afterwards, tilting cryo-stage from 45 ° to 15 ° with −5 ° step to calculate the eucentric position, which is in purpose of keeping the electron beam and ion beam focused on the same position. The *in situ* gas injection system (GIS, FEI) coating with Pt is necessary to improve sample conductivity and reduce curtaining artifacts during FIB milling (59) with the GIS coating time of 6 s. After the GIS coating, the samples usually have a more extensive charging, normally the sputtering coating is useful to improve the overall conductivity of the sample (20~30 mA, 10 Pa and 15 s). The lamellas were milled using the Ga^+^ ion beam at 30 kV under a shallow of 14~20 ° angle. Rough milling was performed with a rectangular pattern (8 × 3 ×10 μm) and 0.5 nA beam current, followed by sequentially lowered beam currents of 0.3 nA, 0.1 nA and 50 pA during the thinning and cleaning steps. During the milling, the sample was also imaged using the CCD and T2 camera at 2 kV and 10 pA (40). The rough milling is usually stopped at the lamella depth of 800 nm. When all the lamellas were rough milled, the final polishing can be done with the current of 30 pA or 10 pA. The desired lamella thickness is typically around 200 nm and around 5~7 lamellas were milled one day. Please note that contamination will grow on lamella eventually, so fast polishing of the lamella ensures it within the desired final thickness. Around 40 lamellas of MCP and 25 lamellas of WCP were milled. The milled grids were stored in sealed boxes in liquid nitrogen for cryo-electron tomography imaging.

### Cryo-electron tomography

The milled grids were transferred to a 300 −kV titan krios electron microscope (FEI, Thermo Fisher) equipped with an energy filter and a heated phase plate holder (60). Images were recorded at 26,000 × magnification with pixel size of 5.4 Å. SerialEM (61) was used to collect tilt series at −6 μm defocus, 4 s record time and a cumulative dose of ~80 e^−^/Å (62). The image stacks were collected at a tilt angles between −38 ° and +64 ° with a tilt step of 3 ° (considering the cryo-FIB lamella tilt angle of 20 °). The dataset for this study consisted of 22 tomograms from 40 lamellas for MCP and 24 tomograms from 25 lamellas for WCP.

### Reconstruction and segmentation

The FIB-SEM stack of WCP and MCP contains 1199 and 1481 images, respectively. The image sequences of each cell were picked manually and were then aligned and segmented with Amira 6.5 software (Thermo Fisher). After which, the segmented files were smoothed with UCSF Chimera (63). Each tomogram stack contains 35 images, which were first aligned using Motioncorr (64) and were then aligned using fiducial markers by ETOMO (65, 66) or patch tracking with IMOD software (67). The tomograms were reconstructed by siRT at bin 4 using TOMO3D (68). EMAN 2.2 (69–71) were used for the segmentation. UCSF Chimera (72) software packages were used for surface rendering and smoothing of the segmented structure.

The cellular volume, surface area and diameter were measured with the command of label analysis in Amira 6.5, which calculated the average, minimum and maximum of the volume, surface area and diameter of cellular or organelles. We adopted the average value for the calculation. The cellular equivalent diameter was calculated according to the cellular average volume. The relative surface area is the ratio of surface area to volume.

### Proteomic analysis

On the 3^rd^ day of cultivation, microalgae suspension was harvested separately for WCP and MCP and centrifuged to collect the pellet for label-free proteomics. Proteomic analysis was performed with an EASY-nLC^TM^ 1200 UHPLC system (Thermo Fisher) coupled to an Orbitrap Q Exactive HF-X mass spectrometer (Thermo Fisher) (73). The raw data obtained via mass spectroscopy (MS) was used for protein identification based on the corresponding database (P101SC18062128-nannochloropsis-uniprot. fasta; 16455 sequences). Results were further filtered with Proteome Discoverer 2.2 and a false discovery rate (FDR) test was performed to eliminate peptides and proteins above a 5% FDR. Common function annotations of the remaining proteins were performed based on the COG (http://www.ncbi.nlm.nih.gov/COG/), GO (http://www.geneontology.org) and KEGG (http://www.genome.jp/kegg/) databases. During analysis, the protein with fold change (FD=protein of MCP/WCP) > 0.85 and < 1.25 were filtered out. When calculating the functional enrichment of proteins, average FD of each KEGG pathway was calculated and the pathway was filtered out when its average FD > 0.85 and < 1.25.

### Metabolomics analysis

On the 3^rd^ day of cultivation, microalgae suspension was harvested separately for WCP and MCP for untargeted metabolomics. Cell samples were grounded with liquid nitrogen and metabolites were extracted with methanol and formic acid by well vortex. The supernatant collected was injected into the LC-MS/MS system for UHPLC-MS/MS analyses, which were performed using a Vanquish UHPLC system (Thermo Fisher) coupled with an Orbitrap Q ExactiveTM HF-X mass spectrometer (Thermo Fisher). The raw data generated by UHPLC-MS/MS were processed using the Compound Discoverer 3.1 (CD 3.1, Thermo Fisher) to perform peak alignment, peak picking, and quantitation for each metabolite. Using KEGG database and HMDB database (https://hmdb.ca/metabolites) to annotate the metabolites. Orthogonal partial least squares discriminant analysis (OPLS-DA) and T-test were performed for filtering. The metabolites with variable importance plot (VIP) > 1 and P-value < 0.05 were considered to be significant differential metabolites. When calculating the functional enrichment of metabolites, average FD of each KEGG pathway was calculated and the pathway was filtered out when its average FD > 0.75 and < 1.25. In the discussion of the results, some subcellular sites of metabolites and metabolic processes were estimated by extracting gene sequences from KEGG and UNIPROT databases (https://www.uniprot.org) for enzymes related to the metabolic process (especially enzymes related to significant differential metabolites) and performing subcellular localization prediction for those enzymes in plant Mploc (74).

## Acknowledgments

This study was supported by the National Key Research and Development Program-China (2016YFB0601003), Zhejiang Provincial Key Research and Development Program-China (2020C04006). We thank the Center of Cryo-Electron Microscopy (CCEM), Zhejiang University for FIB-SEM.

---

**Figure S1.**
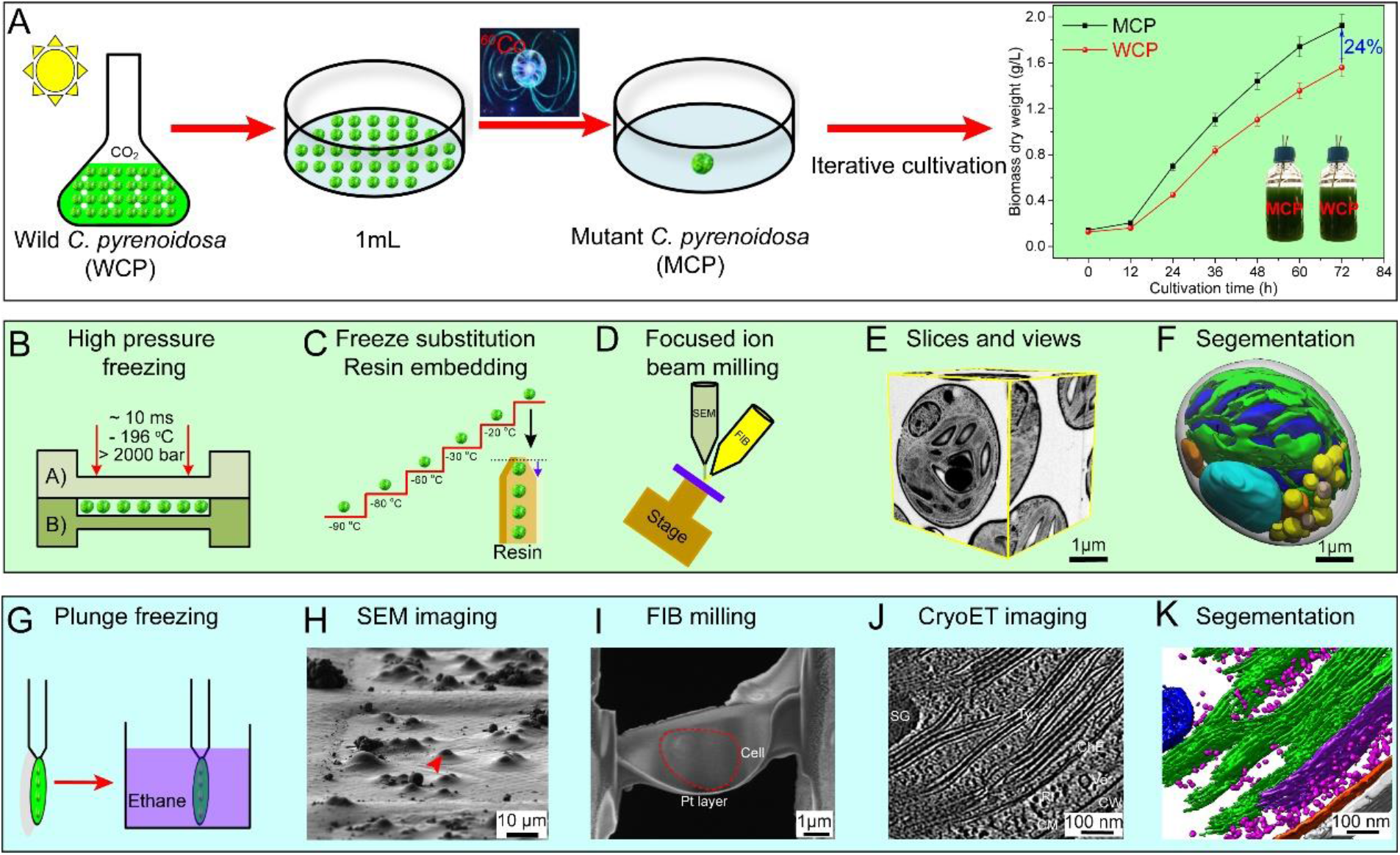
Process of nuclear-irradiation and workflow of subcellular visualization of *C. pyrenoidosa* cells by focused ion beam milling coupled with scanning electron microscopy(FIB-SEM) and cryo-electron tomography technology (Cryo-ET). (A) A high-throughput method for generating gain-of-function mutations in microalgae species using ^60^*Co γ* rays (27, 28). 1 mL of the wild *C. pyrenoidosa* cell suspensions were taken and then radiated under the ^60^*Co γ* rays with the radiation dose of 500 GY and dose-rate of 15 GY/min. The non-directional mutants were then screened to select those with rapid growth rate (15). The mutant *C. pyrenoidosa* cell (MCP) in our experiment have a biomass growth rate 24% higher than that of wildtype (WCP) at the cultivation time of 72 hours. (B-F) Workflow of FIB-SEM. The specimens were frozen at an EM ICE (Leica) high pressure freezing device, where they were imposed to an instant pressure of over 2000 bar within 10 ms, and saved in the liquid nitrogen atmosphere of around −196 °C. The frozen specimen was then substituted in an EM AFS2 (Leica), where the ice crystals were substituted by acetone, and cells were fixed by OsO_4_. The specimen temperature was increased gradually following the procedure in EM AFS2 from liquid nitrogen temperature to room temperature. The resin embedded specimen block was milled by Ga^+^ ion beam and imaged by SEM in the Helios device (Thermo Fisher). The voxel size of the each 2D image is 8 nm and the cell diameter is ~5 μm. In order to obtain the complete dataset, 1199, and 1481 frames of the image stacks were acquired for the WCP and MCP cells, respectively. We obtained clean images with high contrast and low noise for the raw dataset (Video 1,2). The raw images were filtered by the medium 3D filter and manually segmented in Amira software. (G-K) Workflow of cryo-FIB/cryo-ET. (G) Principle of plunge freezing of the *C. pyrenoidosa* cells with back blotting method. (H) The cell imaged with FIB before milling. The frozen specimens were milled by the Ga^+^ ion beam at the liquid nitrogen temperature in the Aquilos device (Thermal Fisher). The target areas were milled by the Ga^+^ ion beam at the tilt angle of 18 °~20 °, and final thickness of the milled lamella was below 200 nm. Totally, 25 lamellas were milled for WCP and 40 lamellas for MCP. (I) Top-view of SEM image of the milled lamella from the same cell shown in (H). The milling direction is from bottom to top. The red-dotted circle is the cell section covered in the lamella. The Pt layer on the bottom of lamella is to protect the sample from damage by the Ga^+^ ion beam. (J, K) A typical tomographic slice of the wild *C. pyrenoidosa* cell and its corresponding 3D segmentation model. The milled grids were then imaged under a 300 kV titan krios electron microscope (Thermal Fisher). It was 24 tomograms collected for WCP and 22 tomograms for MCP. These tomograms were aligned by IMOD software and reconstructed by siRT method at bin 4 using TOMO3D. The image stacks were automatically segmented by machine learning based EMAN2.2 software and smoothed by Chimera software for further investigation. Colors: blue-starch granule, green-thylakoids, magenta-ribosomes, purple-chloroplast envelope, orangered-cell membrane and gray-cell wall.

**Figure S2.**
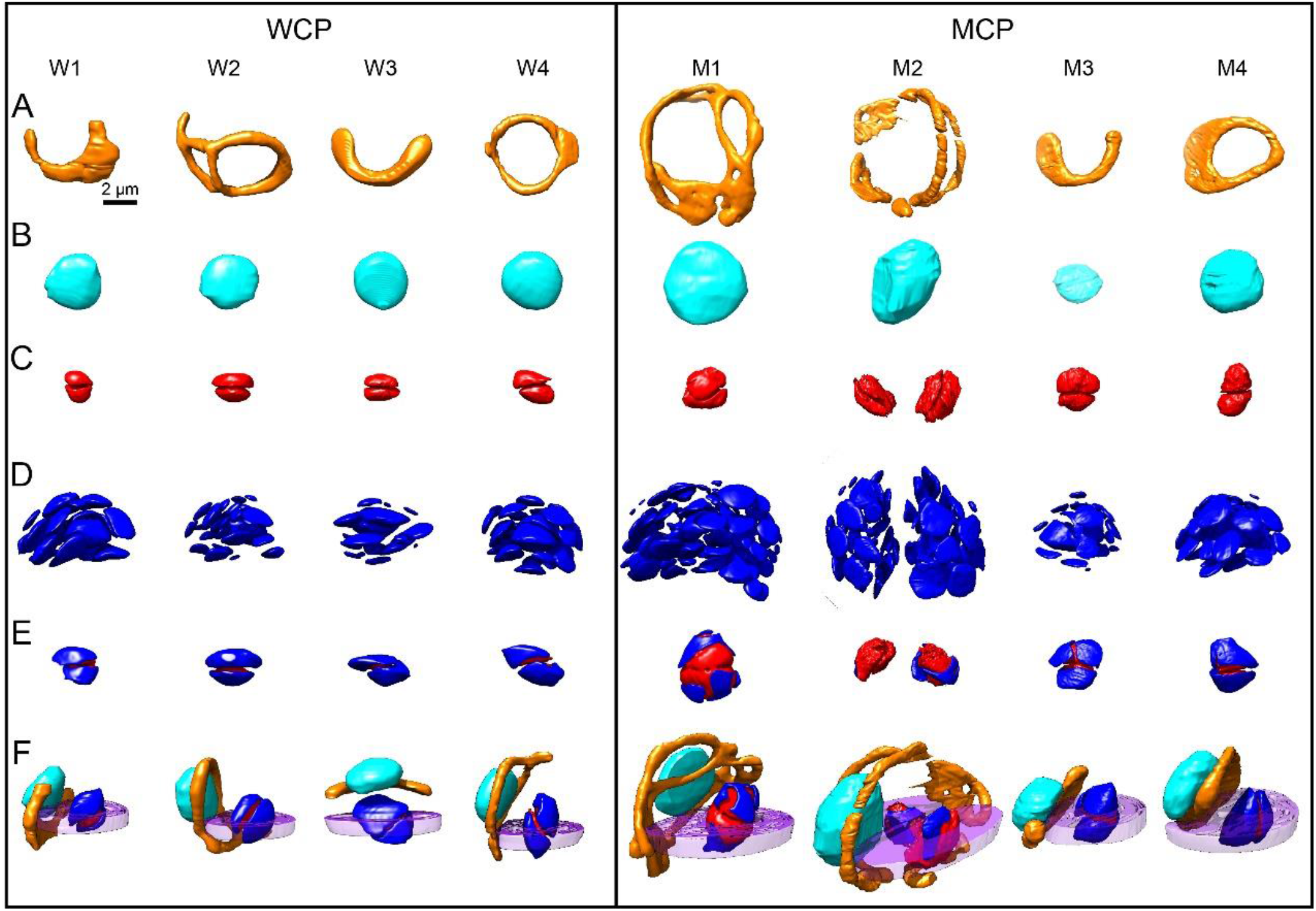
Organelles comparison and special relationship of four wild (WCP, W1-W4) and mutant (MCP, M1-M4) *C. pyrenoidosa* cells, respectively. (A-D) 3D architecture of mitochondria, nucleus, pyrenoid and starch granules in WCP and MCP cells, respectively. (E) The pyrenoid is covered by the starch granules. (F) The spatial relationship of nucleus, mitochondria, chloroplast and pyrenoid. Chloroplast and nucleus are separated by the mitochondria, and the mitochondria is usually prolonged from one side of chloroplast to another side.

**Figure S3.**
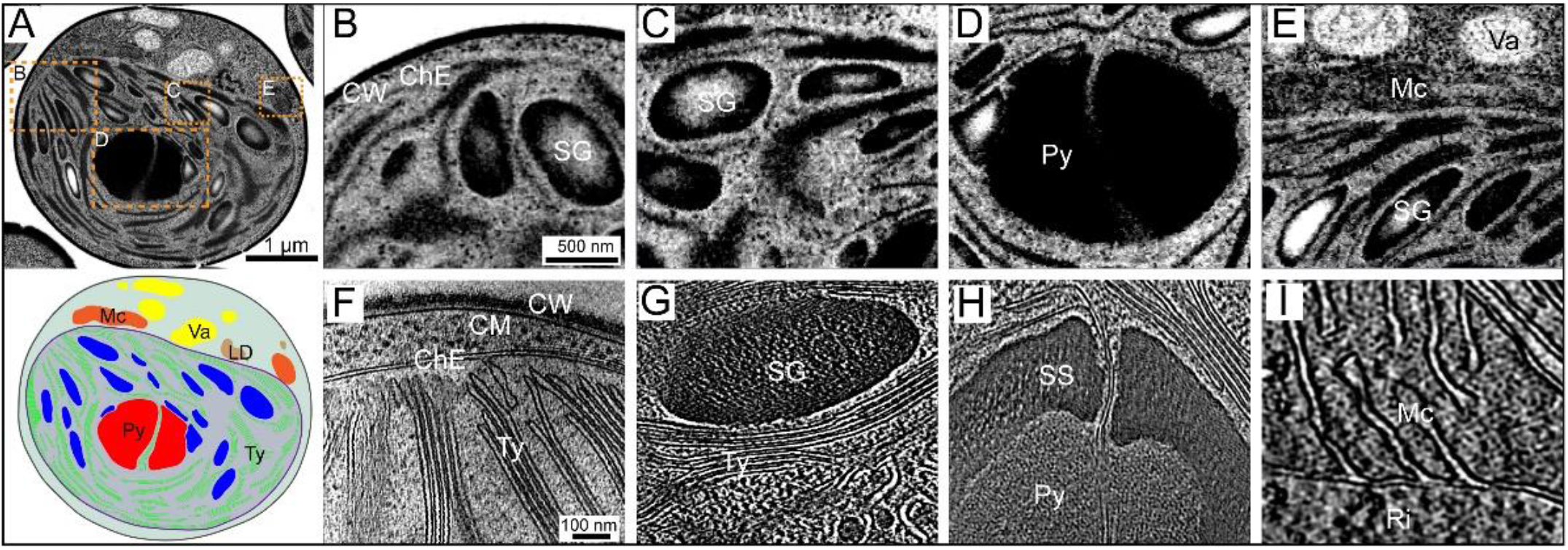
Organelles visualization from FIB-SEM to cryo-ET. (A) A volumetric slice of *C. pyrenoidosa* cell by FIB-SEM and its corresponding 2D diagram. (B, F) The cell wall, chloroplast envelope and thylakoid membrane imaged by FIB-SEM and cryo-ET, respectively. (C, G) The spatial relationship of starch granules and thylakoid membranes imaged by FIB-SEM and cryo-ET, respectively. (D, H) The pyrenoid and starch sheath imaged by FIB-SEM and cryo-ET, respectively. (E, I) The mitochondria section imaged by FIB-SEM and cryo-ET, respectively. CW-Cell wall, ChE-chloroplast envelope, SG-starch granule, Py-pyrenoid, Mc-mitochondria, Va-vacuole, Ty-thylakoid, LD-lipid droplets, SS-starch sheath, Ri-ribosomes.

**Figure S4.**
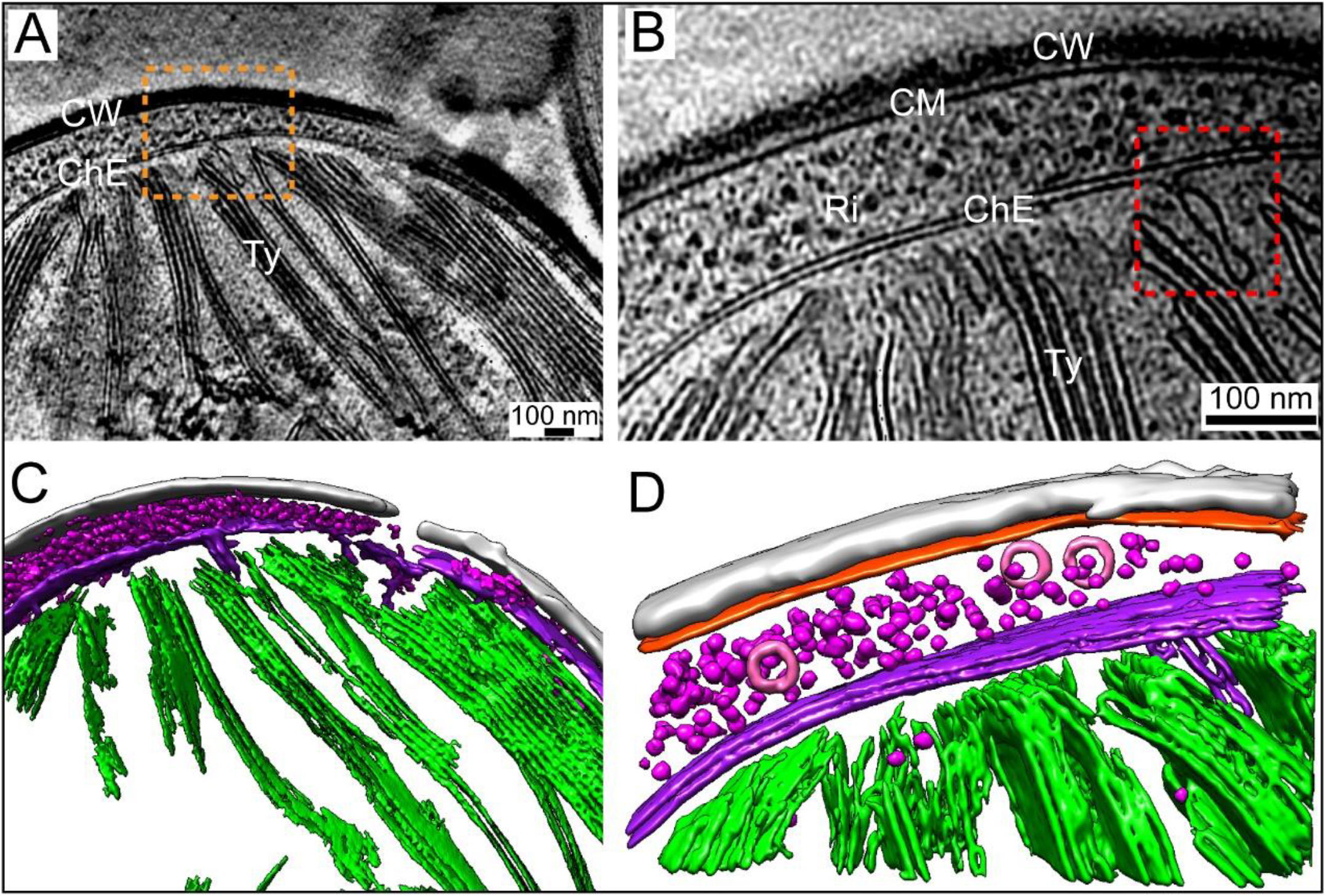
A circular connection channel connecting the chloroplast inner envelope and thylakoid membrane visualized by cryo-ET. (A, C) An overall tomographic slice showing the thylakoid membranes and its corresponding 3D segmentation model. (B, D) Zoom in visualization at the position of orange-dotted box and its corresponding 3D segmentation model. The red-dotted box shows the circular connection channel between chloroplast envelope and thylakoid membrane. Colors: Gray-cell wall (CW), orange red-cell membrane (CM), magenta-ribosomes (Ri), pink-vesicles, purple-chloroplast envelope (ChE), green-thylakoids (Ty).

**Figure S5.**
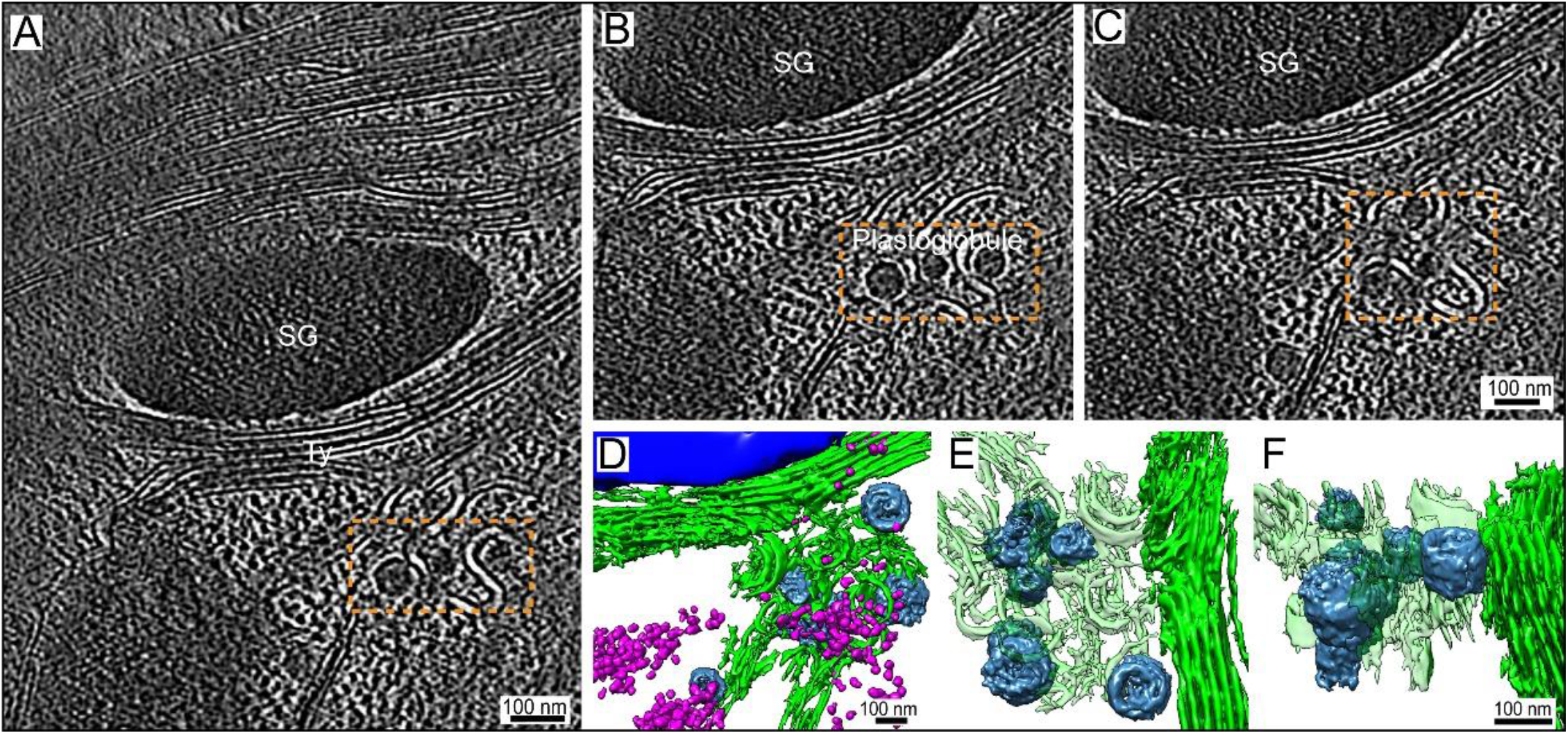
Plastoglobules and starch granules visualized by cryo-ET. (A) A typical tomographic slice showing the starch granules and plastoglobules. (B, C) Two zoom in slices showing the plastoglobules. (D-F) The corresponding 3D segmentation model showing the warped thylakoid membrane and plastoglobules. Colors: Blue-starch granules (SG), steel blue-plastoglobules, green-thylakoids (Ty), magenta-ribosomes.

**Figure S6.**
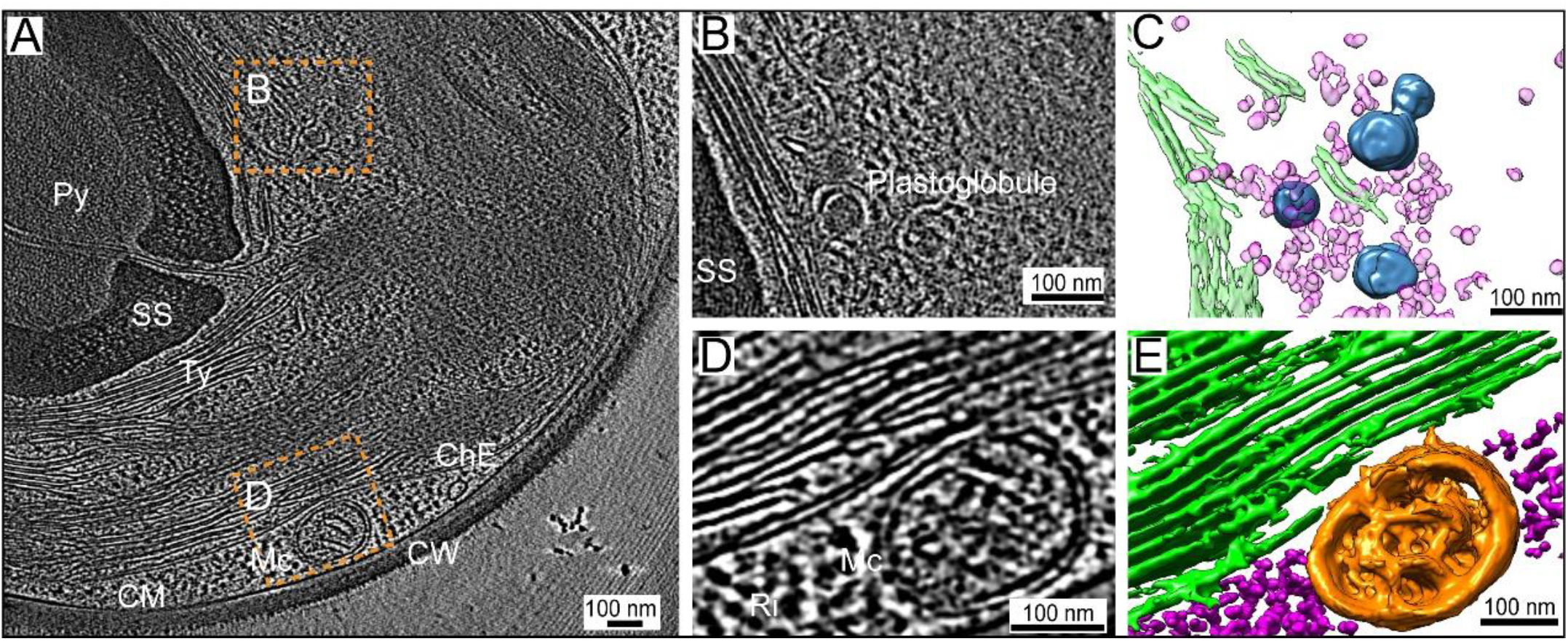
Plastoglobules and mitochondria visualized by cryo-ET. (A) A typical tomographic slice showing the plastoglobules (steel blue) and mitochondria. (B, C) Plastoglobules identified alongside the starch sheath and its corresponding 3D segmentation model. (D, E) A mitochondria section visualized outside the chloroplast envelope and its corresponding 3D segmentation model. CW-Cell wall, CM-Cell membrane, ChE-chloroplast envelope, Py-pyrenoid, Mc-mitochondria (orange), Ty-thylakoid (green), SS-starch sheath, Ri-ribosomes (pink).

**Figure S7.**
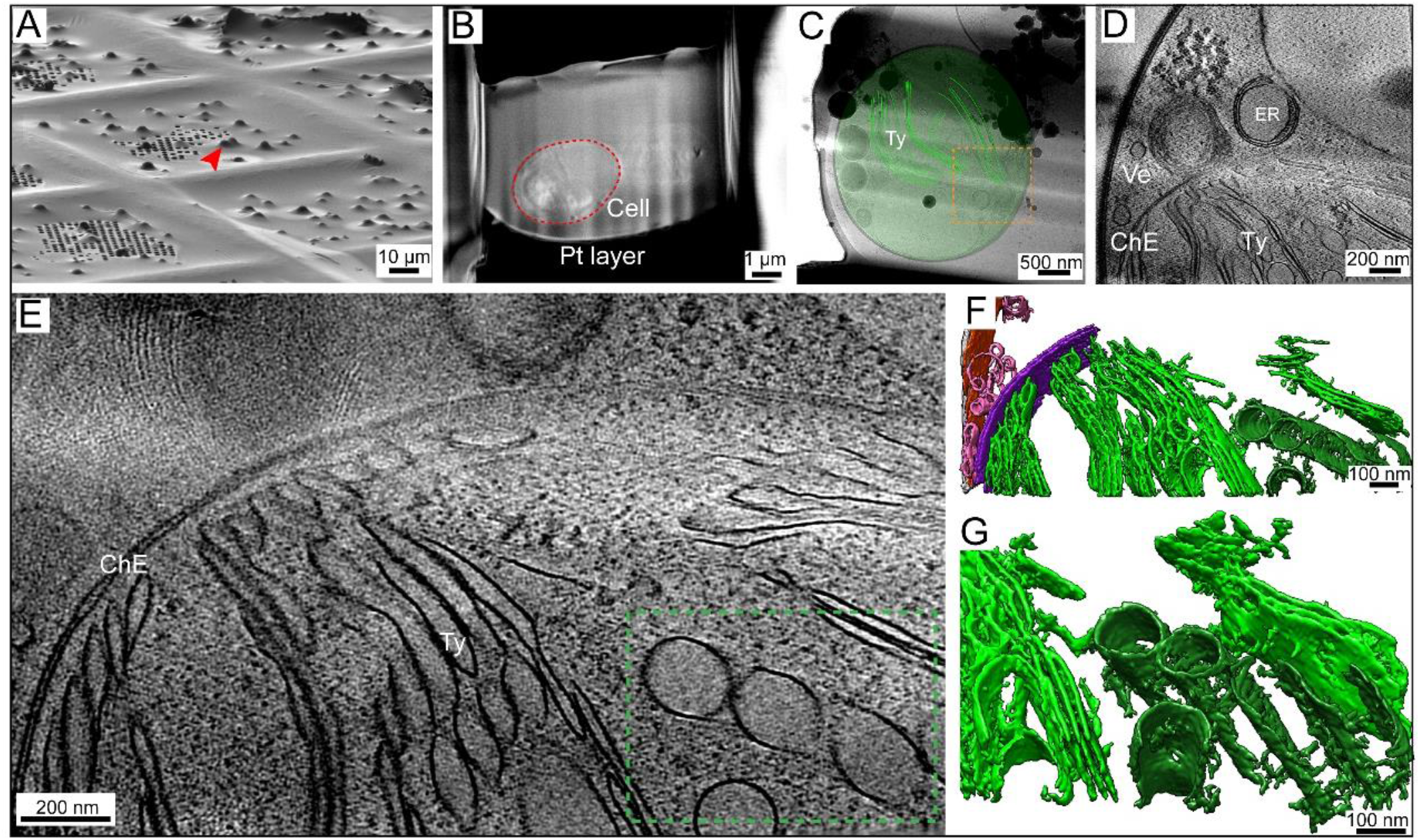
A cryo-FIB milled lamella of mutant *C. pyrenoidosa* cell showing the microtubule-like architecture in the chloroplast matrix. (A) The cell imaged with the FIB before milling at the tilt angle of 18°. (B) Top-view of SEM image of the milled lamella from the same cell shown in (A). (C) TEM high-defocus montaged overview of the same lamella in (B). (D) A typical tomographic slice visualized in the orange-dotted box of (C). Thylakoids, chloroplast envelope, vesicles and endoplasmic reticulum (ER) were visualized. (E) Zoom in visualization of (D) showing the inside structure of chloroplast. (F-G) The corresponding 3D segmentation model. Colors: green–thylakoids (Ty), forest green– microtubules, purple–chloroplast envelope (ChE), pink–vesicles (Ve).

**Figure S8.**
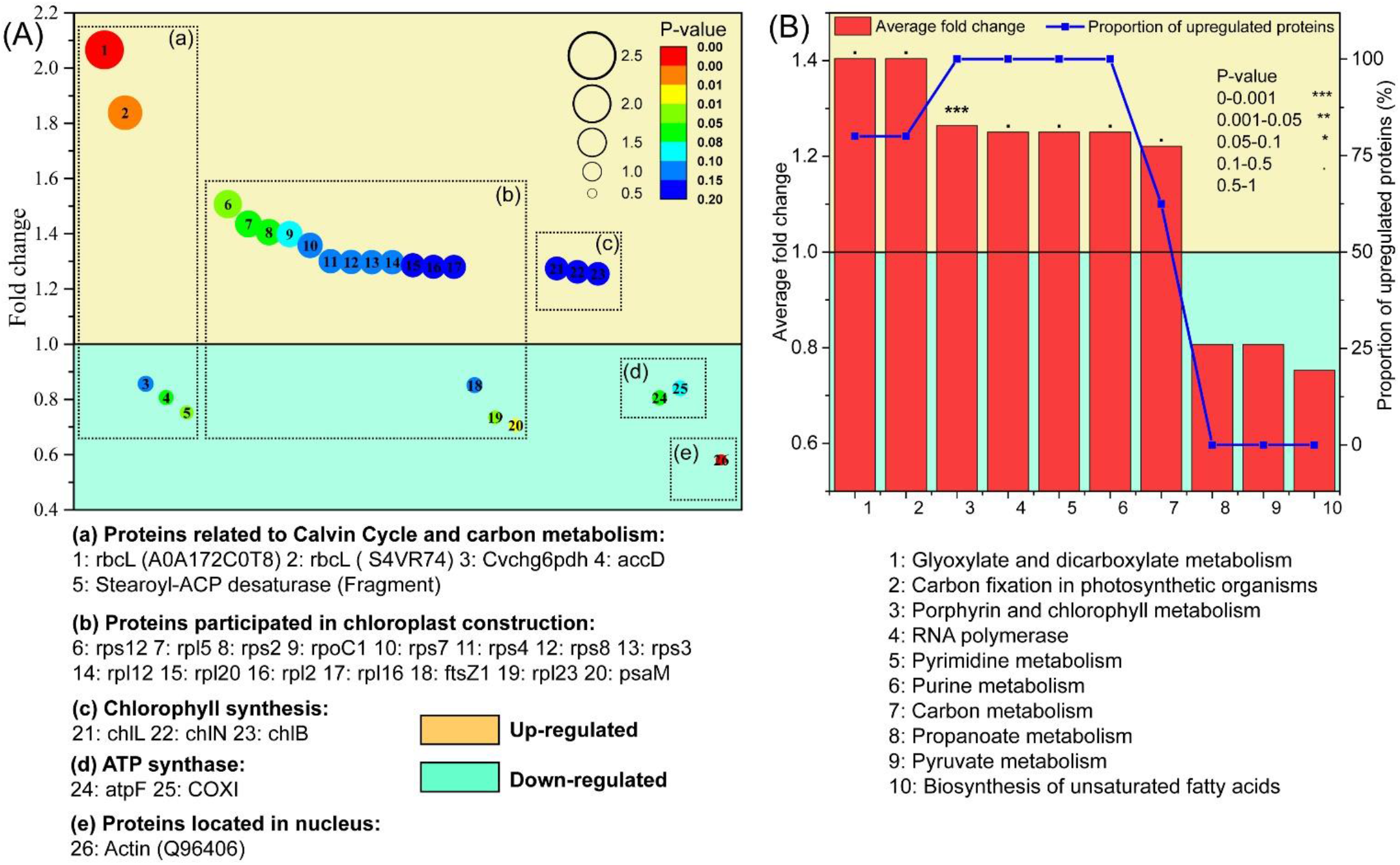
Analysis of proteomics with mutant (MCP) and wild type (MCP) of *C. pyrenoidosa* cell. (A) Abundance change of proteins with some specific functions caused by nucleus mutation; (B) enrichment of protein abundance differences caused by nucleus mutation classified on KEGG pathway (according to KEGG database).

**Figure S9.**
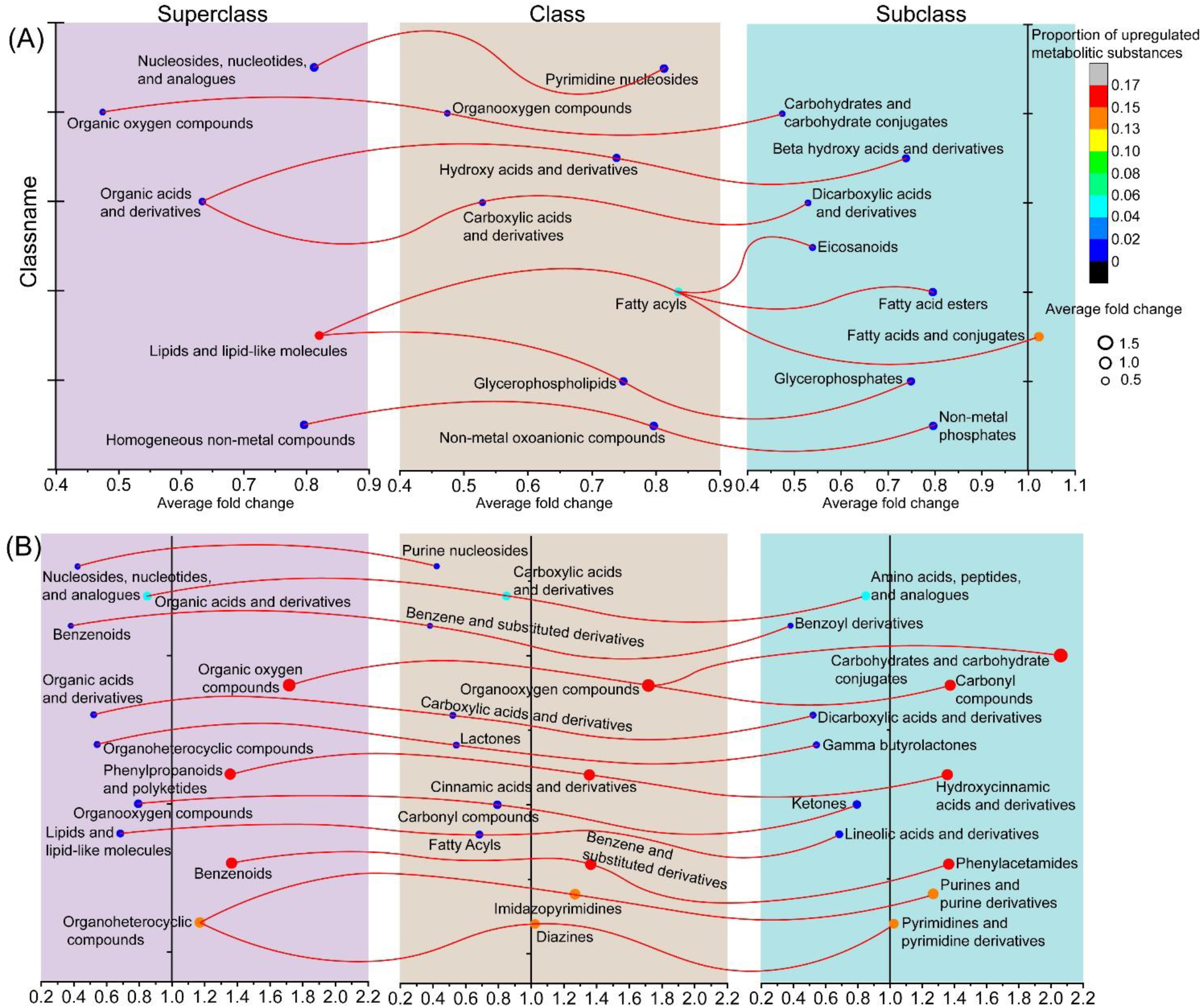
Classification based on HMDB (Human Metabolome Database) class of significant different metabolites of *C. pyrenoidosa* caused by nucleus mutation. (A) Negative ion mode of metabonomics; (B) positive ion mode of metabonomics. Notes: Superclass: the broad categories of metabolites according to HMDB database; Class: the categories of metabolites according to HMDB database, a subordinate classification of Superclass; Subclass: the categories of metabolites according to HMDB database, a subordinate classification of Class.

**Figure S10.**
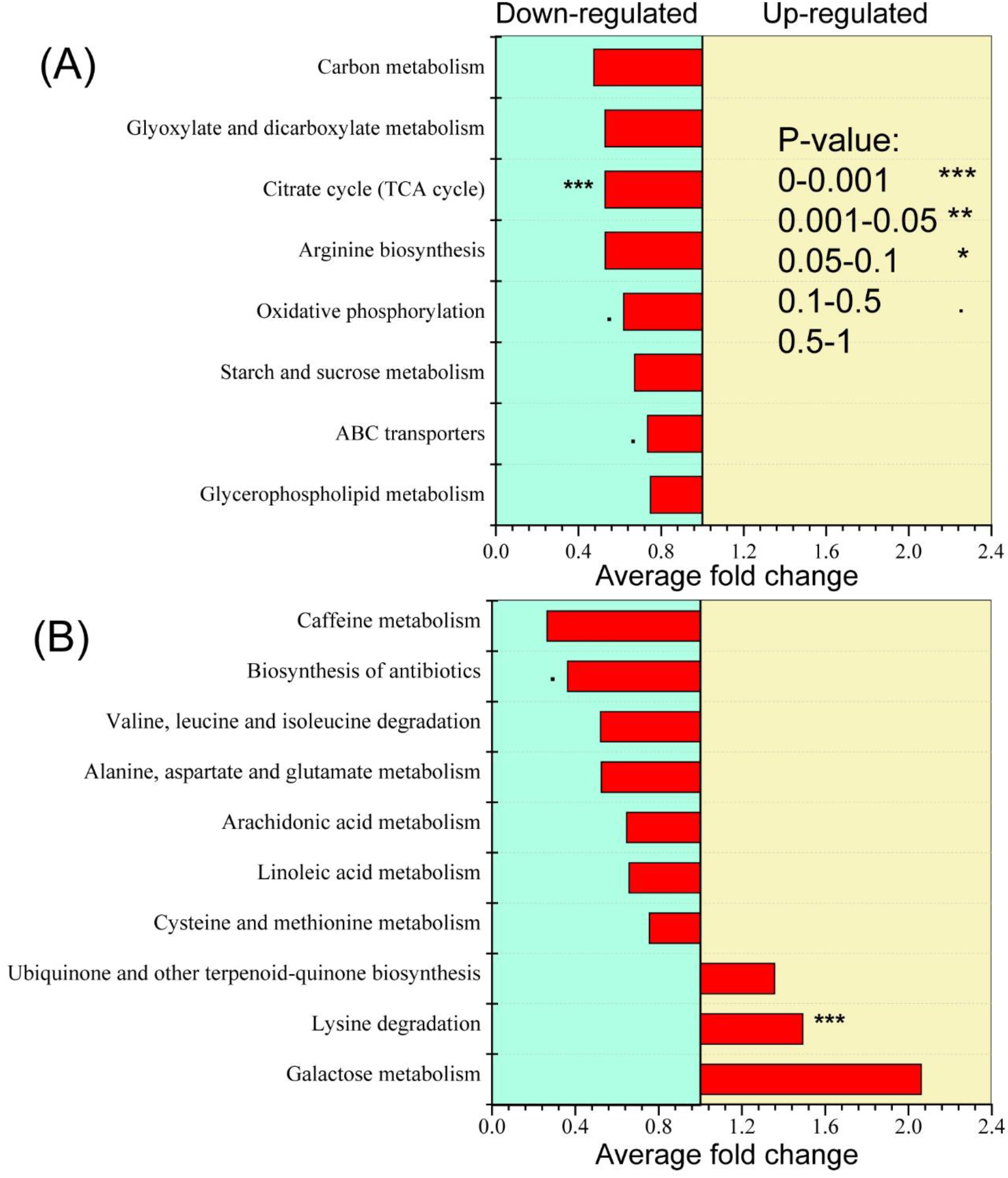
Classification based on KEGG pathway (according to KEGG database) of significant different metabolites of *C. pyrenoidosa* caused by nucleus mutation. (A) Negative ion mode of metabonomics; (B) positive ion mode of metabonomics.

**Figure S11.**
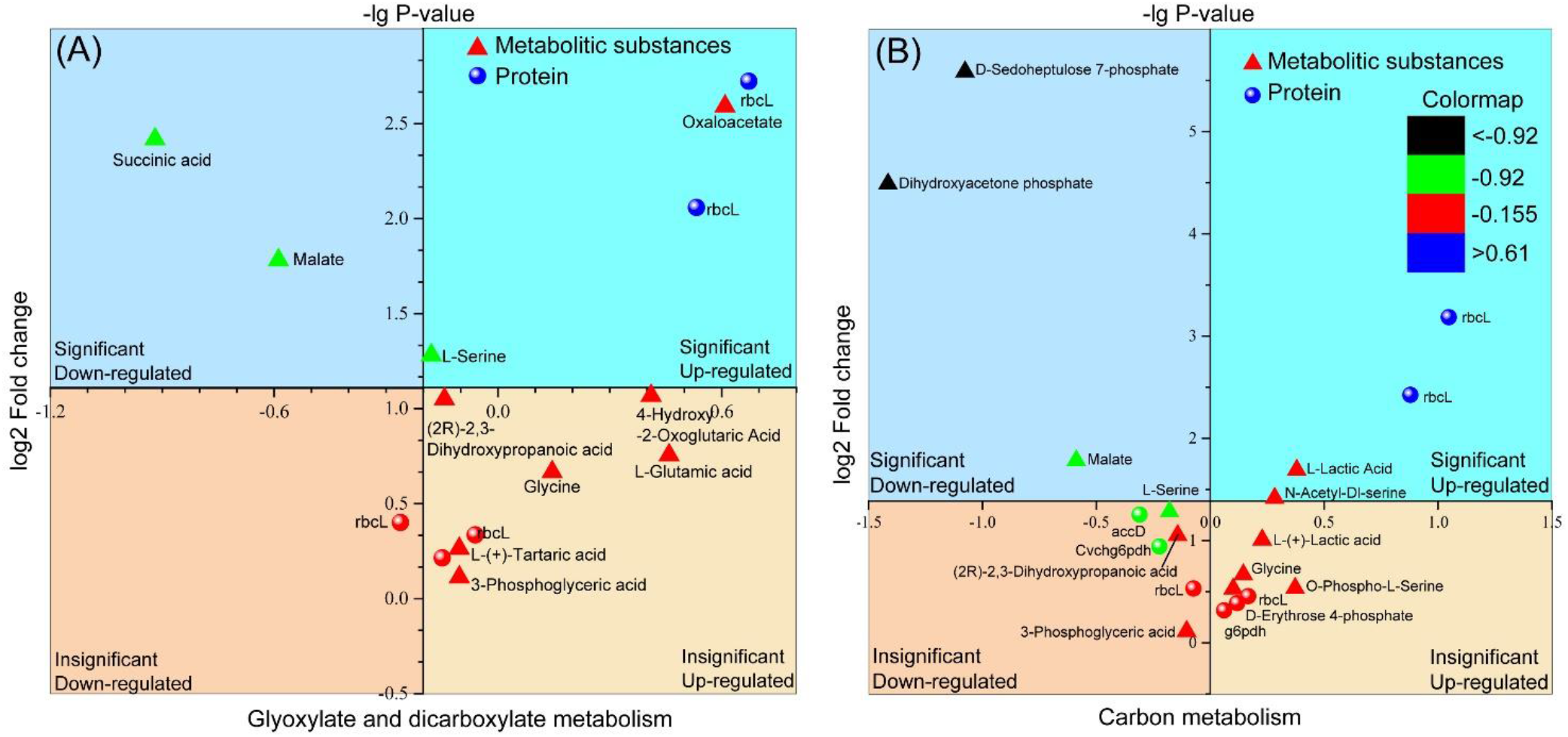
Abundance changes caused by nucleus mutation of proteins and metabolites in the pathways (according to KEGG database) with significant mutation enrichment co-occurring in metabolomics and proteomics. (A) Glyoxylate and dicarboxylate metabolism; (B) Carbon metabolism.

**Table S1.** The volume of the whole cell and different organelles of wild (WCP) and mutant (MCP) *C. pyrenoidosa* cells (μm^3^)

|  |  | Cell | Ch | Ty | SG | CW | Nu | Mc | Py | Va | LD | Cytoplasmic<br>matrix | Chloroplast<br>matrix |
| --- | --- | --- | --- | --- | --- | --- | --- | --- | --- | --- | --- | --- | --- |
| WCP | W1 | 23.97 | 13.72 | 2.74 | 3.92 | 2.11 | 1.44 | 0.75 | 0.35 | 0.34 | 0.11 | 5.50 | 6.71 |
|  | W2 | 27.29 | 13.92 | 1.91 | 1.58 | 1.53 | 1.84 | 0.96 | 0.57 | 0.06 | 0.07 | 8.90 | 9.85 |
|  | W3 | 19.60 | 10.98 | 1.94 | 1.78 | 1.49 | 1.33 | 0.65 | 0.39 | 0.18 | 0.11 | 4.86 | 6.87 |
|  | W4 | 29.30 | 15.51 | 3.21 | 3.26 | 1.98 | 1.70 | 0.73 | 0.52 | 0.30 | 0.07 | 9.00 | 8.52 |
| MCP | M1 | 82.10 | 50.39 | 11.50 | 5.81 | 4.15 | 3.89 | 2.65 | 2.15 | 1.54 | 0.15 | 19.33 | 30.90 |
|  | M2 | 77.10 | 44.30 | 10.76 | 5.72 | 4.69 | 4.15 | 2.30 | 1.12 | 1.34 | 0.12 | 20.21 | 26.70 |
|  | M3 | 24.49 | 13.37 | 3.11 | 1.16 | 1.62 | 1.61 | 0.77 | 0.65 | 0.35 | 0.01 | 6.77 | 8.44 |
|  | M4 | 32.16 | 17.90 | 3.41 | 2.11 | 1.83 | 1.80 | 1.05 | 0.70 | 0.36 | 0.07 | 9.15 | 11.68 |

**Table S2.** The diameter of the cell of wild (WCP) and mutant (MCP) *C. pyrenoidosa* cells (μm) Equivalent diameter: Calculated by the volume of *C. pyrenoidosa* cell

|  |  | Diameter of long | Diameter of short | Equivalent diameter |
| --- | --- | --- | --- | --- |
|  |  | axis (μm) | axis (μm) | (μm) |
| WCP | W1 | 3.9 | 3.3 | 3.6 |
|  | W2 | 4.2 | 3.4 | 3.7 |
|  | W3 | 4.2 | 3.0 | 3.3 |
|  | W4 | 3.9 | 3.4 | 3.8 |
| MCP | M1 | 5.7 | 5.1 | 5.4 |
|  | M2 | 5.6 | 4.9 | 5.3 |
|  | M3 | 3.9 | 3.5 | 3.6 |
|  | M4 | 4.5 | 3.6 | 3.9 |
| Average / WCP |  | **4.1** | **3.3** | **3.6** |
| Average / MCP |  | **4.9** | **4.3** | **4.6** |

**Table S3.** The volume ratio of different organelles to the cells of wild (WCP) and mutant (MCP) *C. pyrenoidosa* cells (%)

|  |  | Ch | Ty | SG | CW | Nu | Mc | Py | Va | LD |
| --- | --- | --- | --- | --- | --- | --- | --- | --- | --- | --- |
| WCP | W1 | 57.2 | 11.4 | 16.4 | 8.8 | 6.0 | 3.1 | 1.5 | 1.4 | 0.5 |
|  | W2 | 51.0 | 7.0 | 5.8 | 5.6 | 6.7 | 3.5 | 2.1 | 0.2 | 0.3 |
|  | W3 | 56.0 | 9.9 | 9.1 | 7.6 | 6.8 | 3.3 | 2.0 | 0.9 | 0.6 |
|  | W4 | 52.9 | 11.0 | 11.1 | 6.8 | 5.8 | 2.5 | 1.8 | 1.0 | 0.2 |
| MCP | M1 | 61.4 | 14.0 | 7.1 | 5.0 | 4.7 | 3.2 | 2.6 | 1.9 | 0.2 |
|  | M2 | 57.5 | 14.0 | 7.4 | 6.1 | 5.4 | 3.0 | 1.5 | 1.7 | 0.2 |
|  | M3 | 54.6 | 12.7 | 4.8 | 6.6 | 6.6 | 3.1 | 2.7 | 1.4 | 0.1 |
|  | M4 | 55.7 | 10.6 | 6.6 | 5.7 | 5.6 | 3.3 | 2.2 | 1.1 | 0.2 |

**Table S4.**
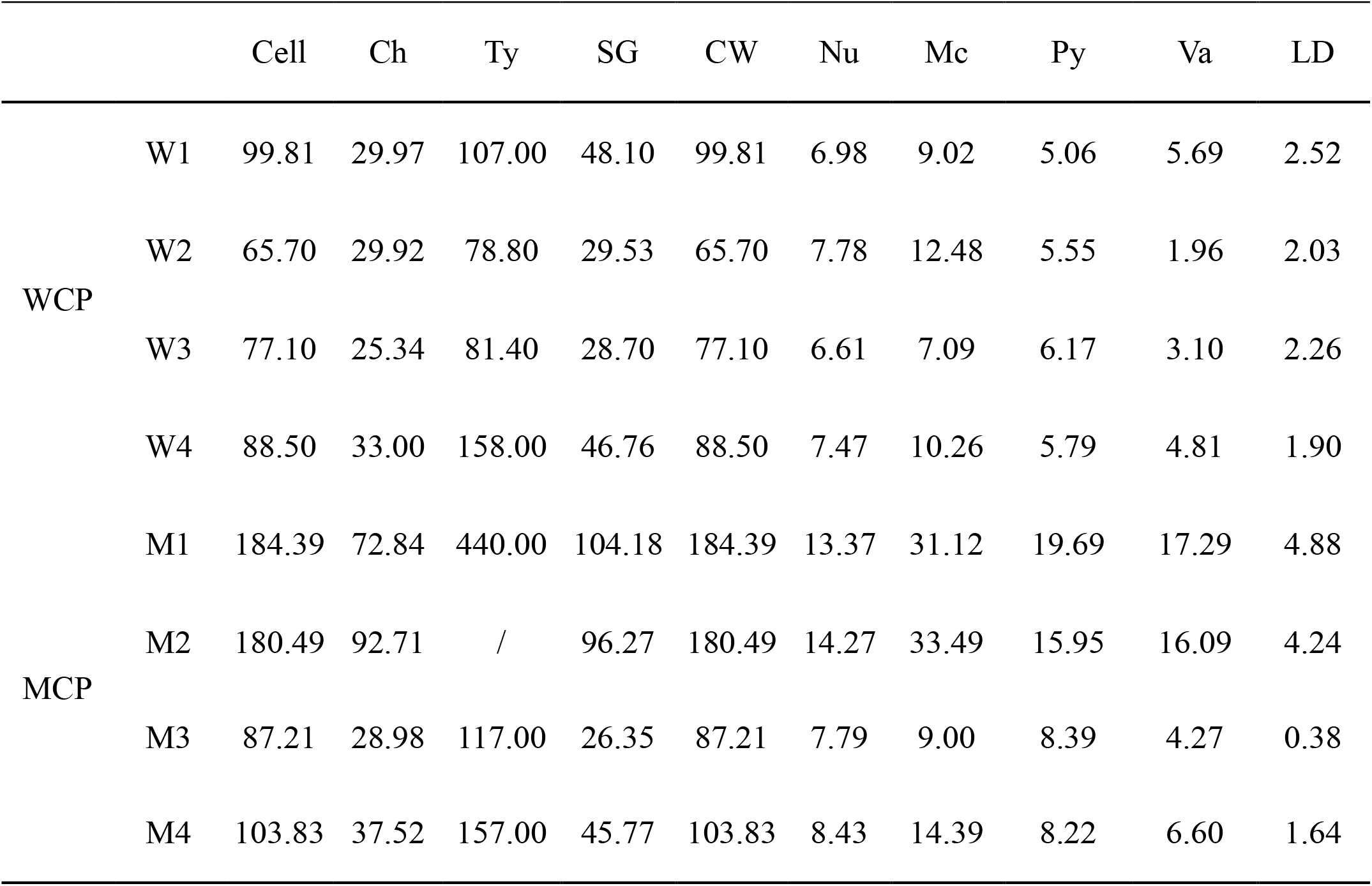
The surface area of the whole cell and different organelles of wild (WCP) and mutant (MCP) *C. pyrenoidosa* cells (μm^2^)

**Table S5.** Relative surface area of the whole cell and different organelles of wild (WCP) and mutant (MCP) *C. pyrenoidosa* cells. Relative surface area: The ratio of surface area to volume of cell or organelles.

|  |  | Cell | Ch | Ty | SG | Nu | Mc | Py | Va | LD |
| --- | --- | --- | --- | --- | --- | --- | --- | --- | --- | --- |
| WCP | W1 | 4.2 | 2.2 | 39.1 | 12.3 | 4.8 | 12.0 | 14.5 | 16.7 | 22.7 |
|  | W2 | 2.4 | 2.2 | 41.3 | 18.7 | 4.2 | 13.0 | 9.7 | 31.6 | 28.9 |
|  | W3 | 3.9 | 2.3 | 42.0 | 16.1 | 5.0 | 11.0 | 15.8 | 16.9 | 20.9 |
|  | W4 | 3.0 | 2.1 | 49.2 | 14.4 | 4.4 | 14.0 | 11.0 | 16.0 | 28.2 |
| MCP | M1 | 2.2 | 1.4 | 38.3 | 17.9 | 3.4 | 11.8 | 9.1 | 11.2 | 31.8 |
|  | M2 | 2.3 | 2.1 | / | 16.8 | 3.4 | 14.6 | 14.2 | 12.0 | 36.0 |
|  | M3 | 3.6 | 2.2 | 37.6 | 22.6 | 4.8 | 11.7 | 12.9 | 12.4 | 27.3 |
|  | M4 | 3.2 | 2.1 | 46.0 | 21.7 | 4.7 | 13.7 | 11.7 | 18.6 | 25.2 |

**Table S6.** The volume ratio of different chloroplast organelles to the chloroplast of wild (WCP) and mutant (MCP) *C. pyrenoidosa* cells (%)

|  |  | Ty | SG | Py | Chloroplast matrix |
| --- | --- | --- | --- | --- | --- |
| WCP | W1 | 20.0 | 28.6 | 2.6 | 48.9 |
|  | W2 | 13.7 | 11.4 | 4.1 | 70.8 |
|  | W3 | 17.7 | 16.2 | 3.6 | 62.6 |
|  | W4 | 20.7 | 21.0 | 3.4 | 54.9 |
| MCP | M1 | 22.8 | 11.5 | 4.3 | 61.3 |
|  | M2 | 24.3 | 12.9 | 2.5 | 60.3 |
|  | M3 | 23.3 | 8.7 | 4.9 | 63.2 |
|  | M4 | 19.0 | 11.8 | 3.9 | 65.3 |

**Table S7.** Measurement of membrane width and gap of thylakoid of wild (WCP) and mutant (MCP) *C. pyrenoidosa* cells (nm)

| Item | Membrane depth |  | Membrane gap |  | Item | Membrane depth |  | Membrane gap |  |
| --- | --- | --- | --- | --- | --- | --- | --- | --- | --- |
|  | MCP | WCP | MCP | WCP |  | MCP | WCP | MCP | WCP |
| 1 | 15.531 | 15.367 | 7.115 | 10.483 | 51 | 14.693 | 9.654 | 7.987 | 9.958 |
| 2 | 13.987 | 17.471 | 7.115 | 9.838 | 52 | 16.846 | 13.913 | 6.477 | 9.341 |
| 3 | 13.913 | 14.774 | 7.717 | 11.237 | 53 | 13.422 | 12.786 | 5.659 | 10.596 |
| 4 | 16.597 | 19.864 | 8.185 | 9.546 | 54 | 15.626 | 10.914 | 7.5 | 10.873 |
| 5 | 16.16 | 17.437 | 8.628 | 10.296 | 55 | 13.199 | 10.804 | 7.979 | 9.325 |
| 6 | 17.837 | 13.187 | 5.148 | 9.293 | 56 | 16.811 | 13.322 | 7.221 | 10.033 |
| 7 | 13.805 | 13.478 | 9.716 | 8.33 | 57 | 18.65 | 11.25 | 6.558 | 10.763 |
| 8 | 15.982 | 11.05 | 8.489 | 8.185 | 58 | 18.336 | 9.762 | 8.248 | 11.678 |
| 9 | 15.193 | 10.497 | 8.799 | 12.117 | 59 | 19.286 | 13.21 | 7.465 | 11.355 |
| 10 | 18.522 | 11.563 | 8.628 | 10.062 | 60 | 20.301 | 10.596 | 7.734 | 10.982 |
| 11 | 18.973 | 13.478 | 8.628 | 11.25 | 61 | 19.006 | 15.444 | 7.465 | 12.739 |
| 12 | 16.155 | 8.33 | 8.185 | 6.988 | 62 | 19.293 | 17.411 | 8.232 | 13.017 |
| 13 | 16.97 | 13.805 | 7.756 | 9.716 | 63 | 21.254 | 12.202 | 7.734 | 10.914 |
| 14 | 17.97 | 9.838 | 8.95 | 11.197 | 64 | 18.066 | 11.473 | 8.375 | 11.25 |
| 15 | 13.117 | 10.539 | 8.112 | 11.25 | 65 | 18.848 | 16.498 | 6.259 | 9.293 |
| 16 | 13.117 | 10.539 | 9.716 | 12.117 | 66 | 15.891 | 17.403 | 9.007 | 10.832 |
| 17 | 14.127 | 13.686 | 9.389 | 10.033 | 67 | 19.524 | 9.654 | 8.761 | 10.426 |
| 18 | 16.155 | 15.014 | 8.883 | 10.596 | 68 | 18.165 | 15.74 | 8.948 | 9.716 |
| 19 | 20.295 | 15.28 | 8.95 | 11.197 | 69 | 16.597 | 14.693 | 9.459 | 12.372 |
| 20 | 15 | 14.03 | 8.883 | 11.805 | 70 | 18.238 | 13.12 | 10.245 | 10.426 |
| 21 | 18.707 | 12.971 | 8.185 | 10.596 | 71 | 20.389 | 12.739 | 7.491 | 11.25 |
| 22 | 17.295 | 13.913 | 9.389 | 8.628 | 72 | 15.162 | 13.913 | 8.185 | 15.512 |
| 23 | 20.295 | 12.202 | 7.342 | 10.721 | 73 | 19.286 | 19.774 | 8.185 | 11.968 |
| 24 | 19.793 | 11.563 | 8.883 | 9.654 | 74 | 18.016 | 16.225 | 9.082 | 13.555 |
| 25 | 18.407 | 11.563 | 8.112 | 8.883 | 75 | 17.573 | 11.602 | 8.057 | 12.018 |
| 26 | 19.591 | 11.918 | 8.489 | 8.542 | 76 | 18.465 | 15.938 | 9.838 | 12.08 |
| 27 | 15.673 | 13.378 | 8.883 | 11.473 | 77 | 17.632 | 15.863 | 7.813 | 10.832 |
| 28 | 15.571 | 12.971 | 8.95 | 9.277 | 78 | 21.001 | 12.263 | 9.546 | 9.654 |
| 29 | 21.25 | 12.263 | 7.87 | 9.325 | 79 | 20.887 | 15.512 | 7.094 | 9.654 |
| 30 | 18.364 | 15.626 | 7.115 | 9.762 | 80 | 17.615 | 16.846 | 7.581 | 10.033 |
| 31 | 18.636 | 15.014 | 8.185 | 8.33 | 81 | 18.303 | 13.773 | 8.883 | 10.968 |
| 32 | 16.917 | 14.693 | 6.661 | 10.354 | 82 | 22.638 | 13.642 | 7.581 | 9.341 |
| 33 | 16.925 | 13.017 | 9.762 | 12.563 | 83 | 15.554 | 13.945 | 8.542 | 9.341 |
| 34 | 17.95 | 12.739 | 8.348 | 8.748 | 84 | 17.257 | 14.083 | 8.33 | 9.277 |
| 35 | 20.374 | 9.958 | 9.293 | 9.958 | 85 | 17.767 | 10.749 | 7.342 | 10.749 |
| 36 | 19.916 | 12.312 | 7.321 | 8.883 | 86 | 15.91 | 10.832 | 6.222 | 11.036 |
| 37 | 15.91 | 13.913 | 10.062 | 9.341 | 87 | 18.066 | 11.981 | 9.53 | 8.799 |
| 38 | 14.825 | 8.731 | 9.082 | 11.395 | 88 | 16.38 | 12.372 | 7.756 | 9.838 |
| 39 | 16.453 | 12.551 | 8.485 | 10.354 | 89 | 15.289 | 12.879 | 8.883 | 10.368 |
| 40 | 16.28 | 11.563 | 7.321 | 8.799 | 90 | 15.093 | 11.678 | 7.87 | 11.473 |
| 41 | 15.626 | 8.628 | 8.647 | 9.53 | 91 | 18.943 | 11.602 | 8.112 | 10.483 |
| 42 | 17.842 | 13.805 | 9.203 | 11.05 | 92 | 16.498 | 11.968 | 7.756 | 9.838 |
| 43 | 18.016 | 12.018 | 7.674 | 9.838 | 93 | 18.016 | 10.483 | 8.112 | 9.033 |
| 44 | 12.456 | 10.426 | 8.462 | 9.762 | 94 | 18.554 | 10.426 | 6.988 | 11.05 |
| 45 | 15.172 | 12.598 | 8.493 | 10.151 | 95 | 15.289 | 13.511 | 7.87 | 9.033 |
| 46 | 12.739 | 12.971 | 9.487 | 11.05 | 96 | 21.394 | 14.744 | 7.342 | 10.296 |
| 47 | 15.521 | 13.289 | 8.624 | 12.263 | 97 | 18.303 | 13.21 | 7.115 | 10.253 |
| 48 | 18.336 | 12.775 | 9.459 | 8.816 | 98 | 18.919 | 10.426 | 8.542 | 8.542 |
| 49 | 14.845 | 13.945 | 7.979 | 10.596 | 99 | 18.409 | 9.654 | 10.539 | 11.13 |
| 50 | 16.925 | 15.377 | 7.175 | 8.472 | 100 | 17.598 | 10.062 | 7.094 | 10.195 |

## Notes

### Competing Interest Statement

The authors have declared no competing interest.

## References

1. J. Silva et al., “Chlorella” in Nonvitamin and Nonmineral Nutritional Supplements. (2019), 10.1016/b978-0-12-812491-8.00026-6, pp. 187–193.

2. J. Anthony et al., An efficient method for the sequential production of lipid and carotenoids from the Chlorella Growth Factor-extracted biomass of Chlorella vulgaris. Journal of Applied Phycology 30, 2325–2335 (2018).

3. J. Doucha, F. Straka, K. Lívanský, Utilization of flue gas for cultivation of microalgae Chlorella sp.) in an outdoor open th in-layer photobioreactor. Journal of Applied Phycology 17, 403–412 (2005).

4. R. Tu et al., “Utilization of Microalgae Cultivated in Municipal Wastewater for CO 2 Fixation from Power Plant Flue Gas and Lipid Production” in Frontiers in Water-Energy-Nexus—Nature-Based Solutions, Advanced Technologies and Best Practices for Environmental Sustainability. (Springer, 2020), pp. 341–343.

5. L. Xu, X. Cheng, Q. Wang, Enhanced Lipid Production in Chlamydomonas reinhardtii by Co-culturing With Azotobacter chroococcum. Frontiers in plant science 9, 741–741 (2018).

6. H. Yu, J. Kim, C. Lee, Nutrient removal and microalgal biomass production from different anaerobic digestion effluents with Chlorella species. Scientific reports 9, 6123–6123 (2019).

7. A. Patel, L. Matsakas, U. Rova, P. Christakopoulos, A perspective on biotechnological applications of thermophilic microalgae and cyanobacteria. Bioresource Technology 278, 424–434 (2019).

8. K. M. Rahman, “Food and High Value Products from Microalgae: Market Opportunities and Challenges” in Microalgae Biotechnology for Food, Health and High Value Products. (Springer, 2020), pp. 3–27.

9. T. Fernandes, A. Martel, N. Cordeiro, Exploring Pavlova pinguis chemical diversity: a potentially novel source of high value compounds. Scientific Reports 10, 1–11 (2020).

10. D. McGee, L. Archer, G. T. Fleming, E. Gillespie, N. Touzet, Influence of spectral intensity and quality of LED lighting on photoacclimation, carbon allocation and high-value pigments in microalgae. Photosynthesis research 143, 67–80 (2020).

11. R. Katiyar et al., Microalgae: An emerging source of energy based bio-products and a solution for environmental issues. Renewable and Sustainable Energy Reviews 72, 1083–1093 (2017).

12. C. Rivasseau et al., Coccomyxa actinabiotis sp. nov.(Trebouxiophyceae, Chlorophyta), a new green microalga living in the spent fuel cooling pool of a nuclear reactor. Journal of phycology 52, 689–703 (2016).

13. W. Wang et al., Repeated mutagenic effects of 60Co-γ irradiation coupled with high-throughput screening improves lipid accumulation in mutant strains of the microalgae Chlorella pyrenoidosa as a feedstock for bioenergy. Algal research 33, 71–77 (2018).

14. S.-W. Poong et al., Transcriptome sequencing of an Antarctic microalga, Chlorella sp. (Trebouxiophyceae, Chlorophyta) subjected to short-term ultraviolet radiation stress. Journal of Applied Phycology 30, 87–99 (2018).

15. J. Cheng et al., Enhancing growth rate and lipid yield of Chlorella with nuclear irradiation under high salt and CO2 stress. Bioresour Technol 203, 220–227 (2016).

16. J. Cheng, Q. Ye, K. Li, J. Liu, J. Zhou, Removing ethinylestradiol from wastewater by microalgae mutant Chlorella PY-ZU1 with CO2 fixation. Bioresource technology 249, 284–289 (2018).

17. Q. Ye et al., Serial lantern-shaped draft tube enhanced flashing light effect for improving CO2 fixation with microalgae in a gas-lift circumflux column photobioreactor. Bioresource technology 255, 156–162 (2018).

18. F. Chu, J. Cheng, X. Zhang, Q. Ye, J. Zhou, Enhancing lipid production in microalgae Chlorella PY-ZU1 with phosphorus excess and nitrogen starvation under 15% CO2 in a continuous two-step cultivation process. Chemical Engineering Journal 375, 121912 (2019).

19. A. Bhattacharya, M. Mathur, P. Kumar, A. Malik, Potential role of N-acetyl glucosamine in Aspergillus fumigatus-assisted Chlorella pyrenoidosa harvesting. Biotechnol Biofuels 12, 178 (2019).

20. T. Lu et al., Evaluation of the toxic response induced by azoxystrobin in the non-target green alga Chlorella pyrenoidosa. Environ Pollut 234, 379–388 (2018).

21. C. Peng et al., Influence of Speciation of Thorium on Toxic Effects to Green Algae Chlorella pyrenoidosa. Int J Mol Sci 18 (2017).

22. D. A. G. a. D. J. Griffiths, The fine structure of autotrophic and heterotrophic cells of Chlorella vulgaris (Emerson strain). Plant & Cell Physiol. 10, 11–19 (1969).

23. G. Gärtner et al., Мicroscopic investigations (LM, TEM and SEM) and identification ofChlorellaisolate R-06/2 from extreme habitat in Bulgaria with a strong biological activity and resistance to environmental stress factors. Biotechnology & Biotechnological Equipment 29, 536–540 (2015).

24. J. A. Schiel et al., Endocytic membrane fusion and buckling-induced microtubule severing mediate cell abscission. J Cell Sci 124, 1411–1424 (2011).

25. C. Uwizeye et al., In-cell quantitative structural imaging of phytoplankton using 3D electron microscopy. bioRxiv 10.1101/2020.05.19.104166 (2020).

26. J. Arnold et al., Site-Specific Cryo-focused Ion Beam Sample Preparation Guided by 3D Correlative Microscopy. Biophysical journal 110, 860–869 (2016).

27. Y. Huang et al., Transcriptome and key genes expression related to carbon fixation pathways in Chlorella PY-ZU1 cells and their growth under high concentrations of CO 2. Biotechnology for biofuels 10, 181 (2017).

28. J. Cheng et al., Mutate Chlorella sp. by nuclear irradiation to fix high concentrations of CO2. Bioresour Technol 136, 496–501 (2013).

29. P. G. Fabrice Rebeille, Interaction between Chloroplasts and Mitochondria in Microalgae. Plant Physiol 88, 973–975 (1988).

30. Q. Hu et al., Microalgal triacylglycerols as feedstocks for biofuel production: perspectives and advances. Plant J 54, 621–639 (2008).

31. P. Přibyl, V. Cepák, V. Zachleder, Production of lipids and formation and mobilization of lipid bodies in Chlorella vulgaris. Journal of Applied Phycology 25, 545–553 (2012).

32. X. Kang et al., Molecular architecture of fungal cell walls revealed by solid-state NMR. Nat Commun 9, 2747 (2018).

33. J. Nomata, T. Ogawa, M. Kitashima, K. Inoue, Y. Fujita, NB-protein (BchN-BchB) of dark-operative protochlorophyllide reductase is the catalytic component containing oxygen-tolerant Fe-S clusters. FEBS Lett 582, 1346–1350 (2008).

34. A. S. Walker, Russ, W. P., Ranganathan, R. & Schepartz, A., RNA sectors and allosteric function within the ribosome. . PNAS 117, 19879–19887 (2020).

35. J. M. F. M. O’BRIAN, Characterization of a Bradyrhizobium japonicum Ferrochelatase Mutant and Isolation of the hemH Gene. Journal of bacteriology 174, 4223–4229 (1992).

36. D. W. Bollivar, Recent advances in chlorophyll biosynthesis. Photosynth Res 90, 173–194 (2006).

37. V. W. Dzapo, R., Investigation on the enzyme-systems of energy producing metabolism in back muscile and the vitality of pigs. 3. Relations of vitality and carcass composition to enzyme pattern of fatty-acid oxidation, glycolysis-side-path, respiratory enzymes, amount of mitochondria and to the system relation of citrate-cycles glycolyses and respiratory enzymes-glycolyses, respectively. Journal of Animal Breeding and genetics 96, 72–85 (1979).

38. D. J. K. T. L. Tootle, Nuclear Actin: From Discovery to Function. The Anatomical record 301, 1999–2013 (2018).

39. C. M. C. a. S. J. Leevers, Do growth and cell division rates determine cell size in multicellular organisms? Journal of Cell Science 113, 2927–2934 (2000).

40. B. D. Engel` et al., Native architecture of the Chlamydomonas chloroplast revealed by in situ cryo-electron tomography. Elife 4 (2015).

41. A. K. Itakura et al., A Rubisco-binding protein is required for normal pyrenoid number and starch sheath morphology in Chlamydomonas reinhardtii. Proc Natl Acad Sci U S A 116, 18445–18454 (2019).

42. E. R. Freeman et al., The Eukaryotic CO2-Concentrating Organelle Is Liquid-like and Exhibits Dynamic Reorganization. Cell 171, 148–162 (2017).

43. L. C. M. Mackinder et al., A Spatial Interactome Reveals the Protein Organization of the Algal CO 2 -Concentrating Mechanism. Cell 171, 133 (2017).

44. J. D. Rochaix, The Pyrenoid: An Overlooked Organelle Comes out of Age. Cell 171, 28–29 (2017).

45. A. Mukherjee, CO2 Concentration in Chlamydomonas reinhardtii: Effect of the Pyrenoid Starch Sheath. Plant Physiol 182, 1796–1797 (2020).

46. C. Toyokawa, T. Yamano, H. Fukuzawa, Pyrenoid Starch Sheath Is Required for LCIB Localization and the CO2-Concentrating Mechanism in Green Algae. Plant Physiol 182, 1883–1893 (2020).

47. I. Pottosin, S. Shabala, Transport Across Chloroplast Membranes: Optimizing Photosynthesis for Adverse Environmental Conditions. Mol Plant 9, 356–370 (2016).

48. I. Pottosin, O. Dobrovinskaya, Ion Channels in Native Chloroplast Membranes: Challenges and Potential for Direct Patch-Clamp Studies. Front Physiol 6, 396 (2015).

49. A. Mechela, S. Schwenkert, J. Soll, A brief history of thylakoid biogenesis. Open Biol 9, 180237 (2019).

50. J. R. Austin2nd, E. Frost, P. A. Vidi, F. Kessler, L. A. Staehelin, Plastoglobules are lipoprotein subcompartments of the chloroplast that are permanently coupled to thylakoid membranes and contain biosynthetic enzymes. Plant Cell 18, 1693–1703 (2006).

51. C. Brehelin, F. Kessler, K. J. van Wijk, Plastoglobules: versatile lipoprotein particles in plastids. Trends Plant Sci 12, 260–266 (2007).

52. S. Rottet, C. Besagni, F. Kessler, The role of plastoglobules in thylakoid lipid remodeling during plant development. Biochim Biophys Acta 1847, 889–899 (2015).

53. E. Lindquist, H. Aronsson, Chloroplast vesicle transport. Photosynth Res 138, 361–371 (2018).

54. M. Rutgers, M. Schroda, A role of VIPP1 as a dynamic structure within thylakoid centers as sites of photosystem biogenesis? Plant Signal Behav 8, e27037 (2013).

55. J. D. P. Heaps, Microtubule-like Structures in the Growing Plastids or Chloroplasts of Two Algae. Planta (Berl.) 81, 193–200 (1968).

56. R. Rippka, J. Deruelles, J. B. Waterbury, M. Herdman, R. Y. Stanier, Generic Assignments, Strain Histories and Properties of Pure Cultures of Cyanobacteria. Microbiology 111, 1–61 (1979).

57. K. Bobik, J. R. Dunlap, T. M. Burch-Smith, Tandem high-pressure freezing and quick freeze substitution of plant tissues for transmission electron microscopy. J Vis Exp 10.3791/51844, e51844 (2014).

58. M. Schaffer et al., Optimized cryo-focused ion beam sample preparation aimed at in situ structural studies of membrane proteins. J Struct Biol 197, 73–82 (2017).

59. M. F. Hayles, D. J. Stokes, D. Phifer, K. C. Findlay, A technique for improved focused ion beam milling of cryo-prepared life science specimens. Journal of Microscopy 226, 263–269 (2007).

60. R. Danev, B. Buijsse, M. Khoshouei, J. M. Plitzko, W. Baumeister, Volta potential phase plate for in-focus phase contrast transmission electron microscopy. Proceedings of the National Academy of Sciences 111, 15635–15640 (2014).

61. D. N. Mastronarde, Automated electron microscope tomography using robust prediction of specimen movements. Journal of structural biology 152, 36–51 (2005).

62. H. Terashima, S. Kojima, M. Homma, Flagellar motility in bacteria: structure and function of flagellar motor. International review of cell and molecular biology 270, 39–85 (2008).

63. E. F. Pettersen et al., UCSF Chimera—a visualization system for exploratory research and analysis. Journal of computational chemistry 25, 1605–1612 (2004).

64. X. Li et al., Electron counting and beam-induced motion correction enable near-atomic-resolution single-particle cryo-EM. Nature methods 10, 584 (2013).

65. A. Korinek, F. Beck, W. Baumeister, S. Nickell, J. M. Plitzko, Computer controlled cryo-electron microscopy–TOM2 a software package for high-throughput applications. Journal of structural biology 175, 394–405 (2011).

66. S. Nickell et al., TOM software toolbox: acquisition and analysis for electron tomography. Journal of structural biology 149, 227–234 (2005).

67. J. R. Kremer, D. N. Mastronarde, J. R. McIntosh, Computer visualization of three-dimensional image data using IMOD. Journal of structural biology 116, 71–76 (1996).

68. J.-I. Agulleiro, J.-J. Fernandez, Tomo3D 2.0–exploitation of advanced vector extensions (AVX) for 3D reconstruction. Journal of structural biology 189, 147–152 (2015).

69. M. Chen et al., Convolutional neural networks for automated annotation of cellular cryo-electron tomograms. Nature methods 14, 983 (2017).

70. G. Tang et al., EMAN2: an extensible image processing suite for electron microscopy. Journal of structural biology 157, 38–46 (2007).

71. I. Rees, E. Langley, W. Chiu, S. J. Ludtke, EMEN2: an object oriented database and electronic lab notebook. Microscopy and Microanalysis 19, 1–10 (2013).

72. T. D. Goddard et al., UCSF ChimeraX: Meeting modern challenges in visualization and analysis. Protein Science 27, 14–25 (2018).

73. M. S. R. Elaswarapu, Methods in Molecular Biology. J. M. Walker, Ed., Genomics Protocols: Second Edition (2008), vol. 439.

74. K. C. Chou, H. B. Shen, Cell-PLoc: a package of Web servers for predicting subcellular localization of proteins in various organisms. Nat Protoc 3, 153–162 (2008).

